# Direct Counting of mRNA Copies Inside Individual Lipid Nanoparticles Using In Situ Lysis and Labeling

**DOI:** 10.64898/2026.05.15.725458

**Authors:** Sadie Graves, Martin Jasinski, Erik Olsén, Albert Kamanzi, Yao Zhang, Jerry Leung, Michael Venier-Karzis, Amin Safaeesirat, Pieter R. Cullis, Sabrina R. Leslie

## Abstract

The optimization of mRNA-lipid nanoparticles (mRNA-LNPs) for therapeutic applications is limited in part by the inadequate characterization of mRNA payload heterogeneity. One current challenge is accurately measuring the number of mRNA copies within individual LNPs, where the standard method of intensity-based mRNA number determination is sensitive to fluorescent dye-dye interactions and heterogeneity of mRNA labeling. Here we present a single-particle microscopy method that combines direct counting of the mRNA copies per LNP with LNP size measurements. While confined in microwells, individual mRNA-LNPs are lysed to release their cargo and stained with a dye such that the number of mRNA molecules in each well can be directly counted using fluorescence microscopy. Since the method stains the mRNA cargo *in situ*, it enables characterization of LNPs formulated with therapeutic grade (e.g., unlabeled) mRNA. We applied this approach to two Onpattro®-based LNP formulations prepared using different formulation buffers, where the two formulations had different average mRNA copy number, particle size, and fraction of LNPs lacking mRNA. The ability to directly count the number of mRNA molecules in LNPs establishes a complimentary method to intensity-based mRNA number determination and supports the characterization and screening of clinically relevant LNP formulations.

---

Lipid nanoparticles (LNPs) are a highly effective platform for mRNA delivery to cells and show significant promise for the advancement of cancer immunotherapy, gene editing, and protein replacement.^1–3^ Building on the application of mRNA-LNP vaccines against SARS-CoV-2, there is increasing interest in using LNPs to deliver a wide range of cargoes to treat a variety of diseases, such as melanoma and cystic fibrosis.^1,4^ Beyond traditional cargoes such as siRNA and mRNA, LNPs can successfully encapsulate large cargo molecules, such as CRISPR-Cas9 editing systems and self-amplifying RNA (saRNA), whose properties may advance the therapeutic possibilities of LNPs.^5–7^

Despite the widespread use of LNPs, gaps remain in our understanding of their biophysical properties and their relationship to each other, such as the heterogeneity of size and mRNA payload across LNPs within the same formulation.^8,9^ For particle size, the polydispersity index from dynamic light scattering (DLS) measurements is commonly used to estimate sample heterogeneity.^10^ For mRNA payload, the RiboGreen encapsulation efficiency assay is used to determine the percentage of mRNA that is encapsulated within LNPs at the ensemble level.^11,12^ However, this metric measures neither the fraction of empty particles nor the variation in single-particle payload. ^11–13^ As a result, our understanding of intra-sample heterogeneity and its potential relationship with mRNA-LNP efficacy remains limited.^14^

Existing single-particle methods for estimating the number of mRNA molecules per LNP typically rely on rescaling the average payload intensity to a copy number using the signal from a single labeled mRNA molecule, assuming a linear proportionality between the number of mRNA and the fluorescence intensity.^15–17^ However, this approach assumes that the labeling ratio and fluorescence intensity are consistent for all mRNA molecules (Supporting Information, S4.3, Fig. S5C), ^18,19^ ignoring potential for self-quenching^19,20^ and fluorescence resonance energy transfer (FRET).^15,16^ Therefore, unless the linearity is carefully validated, there is an inherent uncertainty of such intensity-based mRNA number estimates. Furthermore, this approach is not always applicable to therapeutic LNPs, as the use of labeled RNA can alter *in vivo* performance. ^21^ In some cases, this can be overcome by using membrane permeable dyes that specifically bind to mRNA.^22–24^ However, relating the fluorescence signal to a mRNA number relies on the assumption that the labeling is homogeneous for all mRNA molecules in the sample, where it has been shown that some membrane permeable dyes label mRNA heterogeneously depending on where the mRNA is located in the LNP.^25^ Therefore, there is a need for complimentary approaches to count the number of mRNA copies in individual LNPs.

To address the need for improved characterization of the mRNA copy number for both labeled and unlabeled cargo, we herein introduce a method to directly count the number of mRNA molecules per LNP at the single-particle level using Convex Lens-induced Confinement (CLiC) microscopy.^16,26,27^ To achieve this, we lyse confined LNPs and stain their mRNA cargo, allowing for single-molecule counting of confined mRNA molecules using fluorescence imaging. Combined with pre-lysis size measurements obtained from single-particle tracking, this method allows us to correlate particle size and mRNA copy number at the single-particle level and investigate a relationship that is not yet well understood.^14,15,28^

## Experimental Section

### Preparation of the lipid nanoparticles

In this study, two different Onpattro®-based LNP formulations were formed by mixing a lipid phase with a 25 mM sodium acetate (NaOAc) or 300 mM sodium citrate (NaCit) buffer at pH 4 containing Firefly luciferase mRNA (1921 nt) via a T-junction mixer before being dialyzed overnight (Supporting Information, S3). ^1,29,30^ In brief, the lipid phase consisted of D-Lin-MC3-DMA (MC3), cholesterol, distearoylphosphatidylcholine (DSPC), and 1,2-dimyristoyl-rac-glycero-3-methoxypolyethylene glycol-2000 (PEG-DMG), and one of 1,1-Dioctadecyltetramethylindodicarbocyanine (DiD) or 3,3’-Dioctadecyloxacarbocyanine perchlorate (DiO) at a molar ratio of 50/38.5/10/1.5/1, respectively. DiD was added to formulations containing unlabeled mRNA and DiO was added to formulations containing Cy5-labeled mRNA to ensure spectral separation between the lipid and mRNA channels.

### CLiC measurements

To enable direct counting of unlabeled mRNA cargo, we used a CLiC flow-cell designed for reagent exchange to trap individual LNPs in microwells, thus allowing LNP lysis, staining of released mRNA molecules, and single-molecule counting using fluorescence microscopy (Fig. 1).^31,32^ Briefly, the flow-cell consists of two coverslips with a 30 µm spacer, where the top coverslip contains an array of cylindrical posts (10µm diameter, 20 nm height, 20 µm spacing) and the bottom coverslip contains an array of microwells (5 µm diameter, 500 nm depth, 3 µm spacing). The posts form a nano-slit (20 nm height) above the microwells that is connected to a microchannel (20-30 µm depth, 200 µm width) on the bottom coverslip to allow reagent exchange (Fig. 1A).

**Figure 1:**
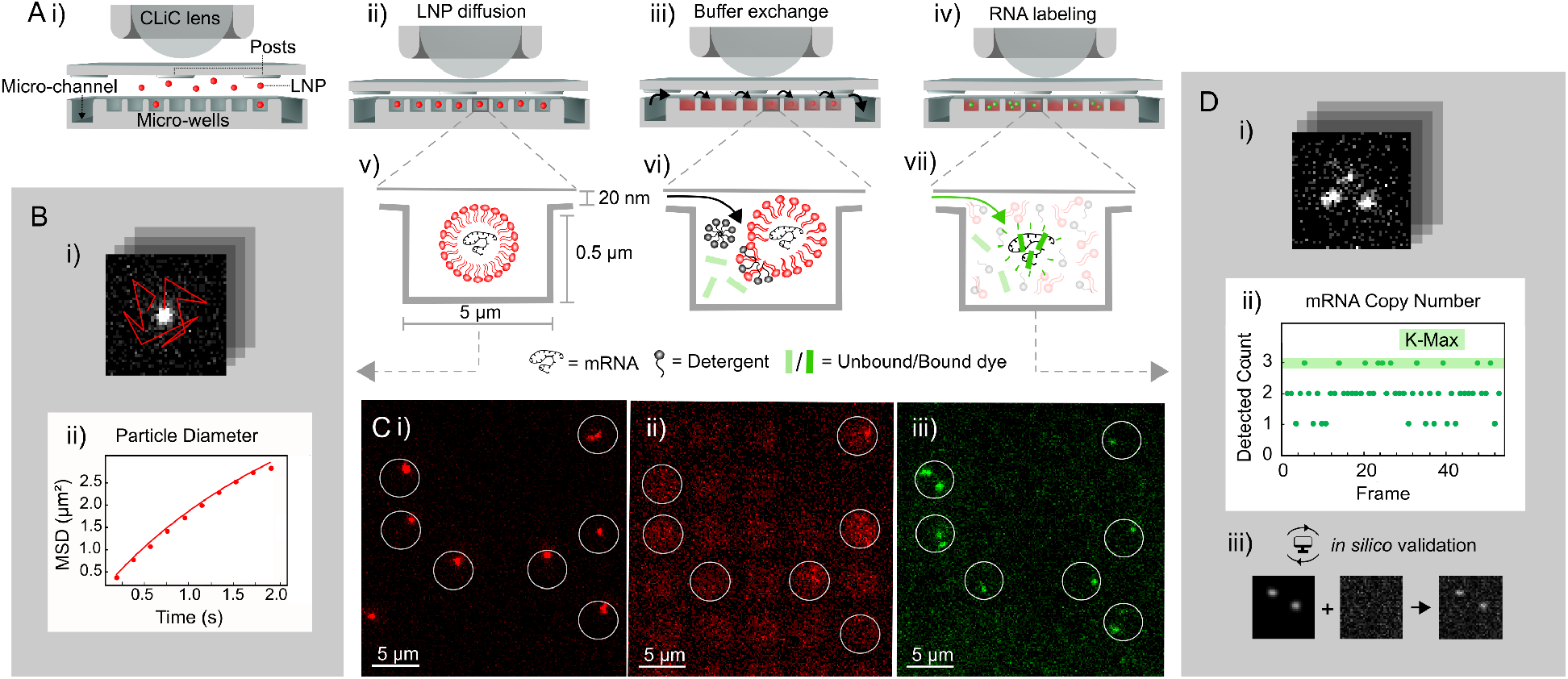
Schematics of the experimental method for LNP lysis and mRNA counting. (A)Schematic of Convex Lens-induced Confinement (CLiC) equipped with reagent exchange for (i) LNP confinement, (ii) size measurements, (iii) buffer exchange to induce LNP lysis, and (iv) RNA labeling. (v) The LNPs are confined in cylindrical microwells with 5 µm diameter and 0.5 µm height, where the 20 nm gap allows (vi) LNP lysis using Triton X-100 and (vii) mRNA staining using RiboGreen while ensuring that the mRNA remains confined in the well. (B) Analysis pipeline for (i) LNP trajectory data to extract particle size via (ii) mean squared displacement (MSD) analysis. (C) Sample images of (i) LNPs pre-lysis, (ii) lysed LNPs, and (iii) post-lysis labeled mRNA. The pre-lysis imaging enables detection of wells containing a single LNP, where only wells with a single LNP are included in the subsequent mRNA counting. (D) Analysis pipeline for (i) mRNA imaging data to determine mRNA copy number from (ii) frame-by-frame counts, as validated with (iii) single-molecule simulations (Supporting Information, S6). The variation in mRNA count between different frames comes from temporary overlapping mRNA molecules.

Before flow-cell assembly, both coverslips were functionalized with 5 kDa polyethylene glycol (PEG) to minimize LNP interactions with flow-cell surfaces, as previously described (Supporting Information, S1.1). ^16,26,31^ Once in-jected into the flow-cell, the LNPs were confined to the microwells by lowering the CLiC pusher lens. The confined LNPs were then imaged for 130 frames to estimate their diffusivity and hence hydrodynamic radii (Fig. 1B and Supporting Information, S2.5), where the ensemble-average result of particle sizing using CLiC has been shown to be in good agreement with other methods.^16^ To minimize the occurrence of multi-particle wells, the bulk dilution of the LNP sample is chosen such that 10− 20% of all wells meet the single-particle criteria, which by Poisson statistics implies that wells containing multiple LNPs occur at a frequency of around 1%. Moreover, the pre-lysis imaging enables detection of wells containing a single LNP using the same detection and counting analysis as for the post-lysis mRNA counting, where only wells with a median particle count of one are included in the subsequent mRNA counting. Next, a mixture of Triton® X-100 (0.2% w/w) and RiboGreen (1:500 dilution) was introduced into the microwells via microfluidic buffer exchange to lyse the confined LNPs and label their mRNA cargo (Fig. 1C) (Supporting Information, S1.2). Finally, the post-lysis labeled mRNA molecules were imaged for 50 frames to enable direct copy number counting. Data was acquired with a Nikon Ti-E inverted microscope using a 100× oil immersion objective and an exposure time of 20 ms (Supporting Information, S1.3).

### Direct mRNA counting

The mRNA counting in this work is based on the detection of local intensity maxima after LNP lysis in each well, where each local maximum above an integrated intensity threshold corresponds to a single mRNA molecule (Fig. 1D). To improve the signal-to-noise (SNR) ratio during single molecule detection, all microscopy videos were denoised with a U-Net model (Supporting Information, S2.1). The intensity maxima of individual LNPs and mRNA molecules were quantified by fitting a 2D Gaussian to the intensity peak(s) (Supporting Information, S2.2 & S2.4). Direct counting was applied to mRNA molecules labeled with either Cy5 or Ri-boGreen. Cy5-labeled mRNA molecules were covalently labeled prior to the particle formulation step, whereas RiboGreen-labeled mRNA molecules were initially unlabeled during the formulation step but later stained *in situ* during the lysis step. Since the number of visible peaks changes between frames due to positional overlap, the number of molecules per well was calculated using a custom statistical estimator that is based on using the median value from frames with the highest mRNA count (Supporting Information, S2.3).

## Results and Discussion

We applied our direct mRNA counting and particle sizing method to two Onpattro®-based LNP formulations manufactured using either NaOAc or NaCit buffer, where these formulation buffers are known to produce distinct differences in LNP structure detectable by Cryo-TEM (Fig. 2A). In particular, a large sub-population of NaCit LNPs contains structures known as blebs, while the use of NaOAc produces LNPs with a solid oil core. This structural difference likely gives rise to different mRNA payloads, where blebs are hypothesized to enclose the mRNA molecules.^33^ Furthermore, the presence of blebs has been reported to affect functional mRNA delivery in mice.^33^

**Figure 2:**
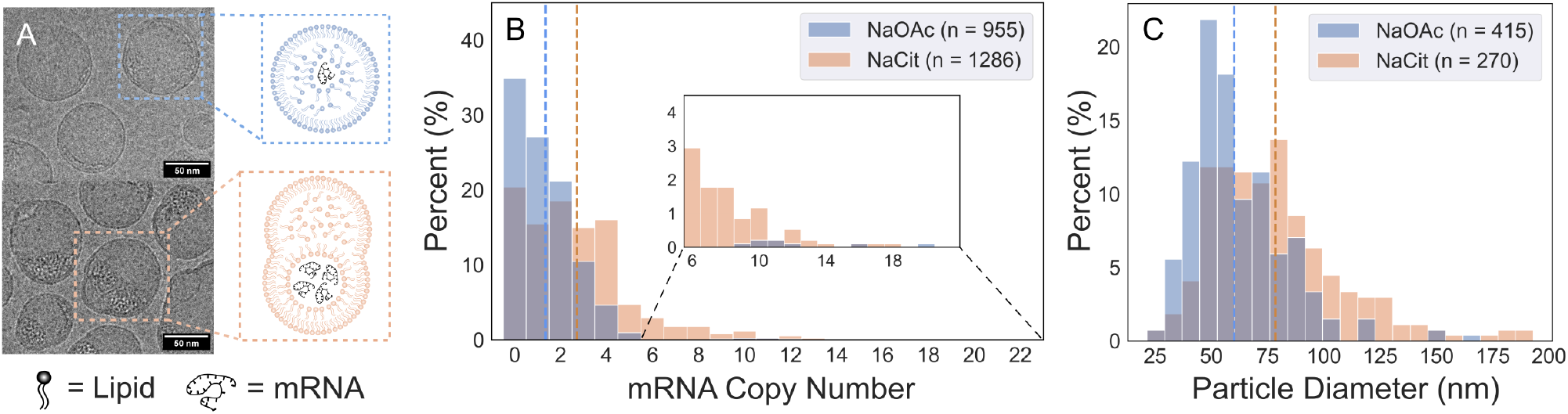
Size and mRNA copy number distributions for LNPs made using two different formulation buffers. (A) Cryo-TEM images of LNPs formulated in NaOAc (top) and NaCit buffer (bottom) with schematics of the structural differences. (B) Distribution of mRNA copy number for LNPs formulated in NaOAc (blue) or NaCit (orange) buffer. The dashed line indicates the mean mRNA copy number (*n*) of the population (*n*_NaOAc_ = 1.3, *n*_NaCit_ = 2.7) and the leftmost bar indicates the empty fraction (*e*), rounded to the nearest percent (*e*_NaOAc_ = 35%, *e*_NaCit_ = 20%). In B, the counts >4 are adjusted using a linear fit on the summed pixel intensities of each well (Supporting Information, S6.3). (C) Distribution of particle diameter for LNPs formulated in NaOAc (blue) or NaCit (orange) buffer. The dashed line indicates the mean diameter (*d*) of the population (*d*_NaOAc_ = 60 nm, *d*_NaCit_ = 78 nm).

For the NaOAc formulation, the mean number of mRNA molecules per particle was 1.3 ± 0.1, with ∼ 35% of LNPs lacking mRNA cargo. In contrast, the NaCit formulation exhibited a higher mean of 2.7 ± 0.1 mRNA molecules per particle, with only ∼20% of particles lacking cargo (Fig. 2B). The NaCit formulation also showed greater polydispersity in mRNA copy number, with a standard deviation of ± 2.3, compared to ± 1.3 for the NaOAc formulation. This increased dispersity arises from a larger subpopulation of LNPs in the NaCit formulation containing >5 mRNA molecules (>13%), whereas such highly loaded particles are rare in the NaOAc formulation (<2%).

In the analysis, a distinction is made between high (≥5) and low (<5) mRNA copy numbers. For LNPs containing ≥5 mRNA molecules, direct mRNA counting becomes unreliable due to spatial overlap (Supporting Information, S2.3). This upper limit of direct counting comes from a compromise between throughput, where larger wells allow accurate (>90%) counting of copy numbers ≥5 (Supporting Information, Section S6.5) but reduces the number of wells within the field of view. Therefore, in microwells containing ≥5 mRNA molecules, the mRNA copy number was estimated by rescaling the post-lysis mRNA fluorescence signal using nearby wells containing <5 mRNA molecules (Supporting Information, S6.3), producing the copy number estimates in the inset of Fig. 2B which are consistent with prior studies. ^16,17,34^ As a result, copy number estimates of ≥5 mRNA are more uncertain than those of <5 mRNA molecules. However, since most LNPs (>75%) contain <5 mRNA, this uncertainty has minimal impact on the estimation of the mean copy number. Moreover, the increased spatial separation of mRNA molecules in the wells post-lysis reduces FRET and self-quenching, enabling a calibration of fluorescence intensity to mRNA copy number for formulations containing labeled mRNA (Supporting Information, S5.2, Fig. S7).

In addition to measuring post-lysis labeled mRNA, the mRNA counting method was also applied to LNPs containing Cy5-labeled mRNA, which were also formulated in either NaOAc or NaCit buffers (Supporting Information, S5.1). The average mRNA copy number and overall loading distribution using labeled and unlabeled mRNA are similar, with the Cy5-labeled mRNA yielding an average copy number of 1.8 for the NaOAc formulation and 2.6 for the NaCit formulation. This indicates a minimal effect on mRNA loading into individual LNPs when using Cy5-labeled mRNA, which is important considering that pre-formulation labeled mRNA is commonly used in both molecular and in vitro studies of mRNA-LNPs.^14,16,34–36^

In addition to the increased mRNA payload, LNPs formulated in NaCit buffer have a larger mean diameter (78 ± 2 nm) than those formulated in NaOAc buffer (60 ± 1 nm) (Fig. 2C). The size distribution of the NaCit formulation is also more polydisperse than that of the NaOAc formulation, as shown by the long tail of the distribution (Fig. 2C). These size measurements are in agreement with DLS control measurements performed on the same samples (Supporting Information, S3.3, Table S1).

To further investigate the relationship be-tween the amount of mRNA cargo and particle size, we compared LNP diameter distributions for each discrete mRNA copy number (Fig. 3). Our results show a weak correlation between diameter and mRNA copy number for the NaOAc formulation (Fig. 3A), as shown by a Pearson correlation coefficient (*r*) of 0.06. In contrast, there is a weak to moderate correlation (*r* = 0.25) between diameter and mRNA copy number for LNPs formulated in NaCit buffer (Fig. 3B). While precise conclusions on the correlation between mRNA copy number and particle diameter are limited by the number of analyzed particles and the limited LNP track length in this study, our findings agree with previous measurements using labeled mRNA where the correlation between mRNA copy number and particle volume was shown to depend on the formulation buffer.^16^ The same study also reported that there is approximately no size difference between NaOAc LNPs with and without mRNA cargo, while such a size difference is present for NaCit LNPs. Thus, although the single-particle track length can be improved using, for example, label-free imaging of the LNPs,^37^ these results suggest that the size and mRNA counting in this work has the precision to observe differences in scaling between different LNP samples. Considering that these LNP formulations have different structures (Fig. 2A), the difference in scaling could potentially be related to structure, where it has been shown for a few different LNP formulations that the occurrence of blebs appears to coincide with a noticeable size difference between loaded and empty LNPs.^16^ However, investigating such a potential relation in the case of using unlabeled mRNA would require single-particle loading and structure information for a large collection of different LNP formulations with different structures, which is beyond the scope of this investigation.

**Figure 3:**
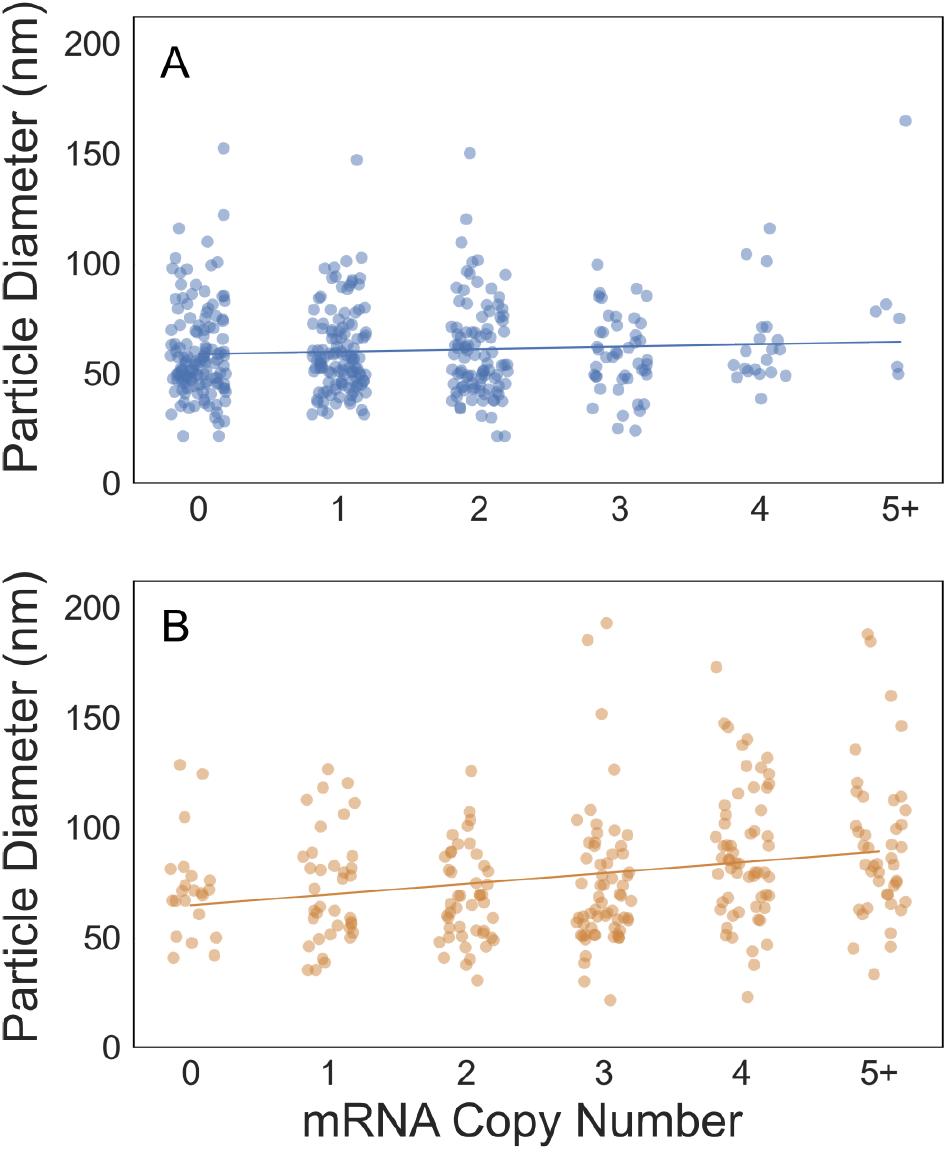
Particle diameter as a function of mRNA copy number. Scatter plots of mRNA copy number as a function of particle diameter for LNPs formulated in (A) NaOAc (blue, n=415) or (B) NaCit (orange, n=270) buffer. The solid line shows the least squares fit. Pearson’s correlation coefficient (*r*) for each LNP formulation was calculated as *r*_NaAOc_ = 0.06 and *r*_NaCit_ = 0.25, with p-values of *p*_NaAOc_ = 0.18 and *p*_NaCit_ <1e-4, respectively.

Overall, our measurements of the average mRNA copy number (1.3-2.7) fall within the approximate range seen in previous studies.^16,34,38,39^ However, there are conflicting reports on whether mRNA copy number is correlated with particle size^38–41^ or if the relationship is more complex.^14,16^ Here, the ratio between the average mRNA copy numbers for the two measured LNP formulations (0.48, Fig. 2B) is similar to the ratio between the average particle volumes (0.45, Fig. 2C), whereas the correlation between size and mRNA copy number within each formulation is limited (Fig. 3A). In the case of a consistent RNA concentration in the LNPs, it is expected that the mRNA copy number scales with particle volume.^16^ Thus, the similarity in ratio between the average particle volume and the average mRNA copy number for the two formulations indicates that the occurrence of blebs does not affect the average mRNA concentration of the LNPs. Moreover, the weak correlation between the mRNA copy number and the particle size at the single-particle level could reflect a Pois-son loading process (Supporting Information, S2.6). Unless the lipid concentration in the LNPs follows the varying mRNA concentration, which would alter the water fraction in the LNPs,^37^ a difference in mRNA copy number without a corresponding volume change would imply a different ratio between lipid and nucleotides. A varying lipid-nucleotide ratio could in turn affect the ratio between ionizable lipids and nucleotides, where a difference in the latter ratio is known to affect LNP function.^14^ Thus, the relationship between particle size and mRNA copy number could potentially be used to obtain information about compositional heterogeneity within an LNP sample, as well as to compare the composition across samples.

## Conclusions

By combining single-particle confinement microscopy with lysis and mRNA staining, we establish here a method for direct counting of the mRNA copy number in individual LNPs. Unlike existing methods, we can measure LNPs loaded with therapeutic grade (e.g., unlabeled) mRNA at the single particle level without relying relating an fluorescence intensity to a mRNA count. For this reason, the direct mRNA counting is a complementary method to the previously used methods for single-particle mRNA counting. Since this method works for both labeled and unlabeled mRNA (Supporting information, S5), we are also able to evaluate the relationship between encapsulated fluorescence intensity and mRNA count.

By combining mRNA counting with single-particle size measurements for two different Onpattro®-based formulations, we show that the relationship between particle size and payload is formulation dependent, however, the average values of particle volume and mRNA copy number scale with each other at the ensemble level. This method also enables the quantification of the fraction of LNPs that do not contain any cargo, which is relevant when determining appropriate dosage in relation to the immunogenicity.^42^ The obtained empty fractions of 20% and 35% are similar to those obtained using labeled mRNA,^16^ which indicates that the empty fraction is not overly sensitive to the use of labeled mRNA.

Although the main focus of this work is direct counting of the mRNA copy number using post-formulation labeled mRNA, the method can also be combined with pre-formulation labeled mRNA, for which we determined a similar mRNA copy number distribution (Supporting Information, S5.1). One interesting possibility of using pre-formulation labeled mRNA is the ability to compare the RNA label intensity of the mRNA-LNP particles prior to lysis, and the direct count following lysis. This has allowed us to evaluate the mRNA number using two different measurement approaches, each of which functions at the single-particle level. Our analysis shows an approximately proportional relationship between these measures (Supporting Information, Fig. S8). However, there is also a significant dispersity when comparing the intensity signal at the single-particle level. This highlights the possibility of combining pre-lysis intensity measurements and direct mRNA counting to explore the heterogeneity of mRNA labeling, which is relevant when evaluating membrane permeable dyes, for example.

Moreover, although the method is optimized to directly count the mRNA copies in the low copy number regime, it also enables count estimates in the larger mRNA number regime, where the distance between mRNA molecules post-lysis minimizes dye-dye interactions. The upper limit for the number of detectable RNA molecules is set by the well size as the direct counting relies on limited spatial overlap between the mRNA molecules (Supporting Information, S6.6). In the case of counting smaller RNA molecules, two criteria must be met. First, it is important that each molecule contains more than one fluorophore to minimize the effects of fluorophore bleaching and blinking. Second, the accessible gap between the PE-Gylated coverslips must be small enough that the RNA molecules cannot easily escape whilst being large enough for the lysis and labeling agents to enter the wells. Since RNA consisting of one thousand nucleotides has a diameter of approximately 20 nm when in solution, ^43^ the current implementation of the mRNA counting method has a lower limit of approximately one thousand nucleotides. Moreover, the low copy number regime is important for therapeutic applications, for which the LNP diameter is generally less than 100 nm and the mRNA is often

>2000 nucleotides.^44^ If the RNA concentration in said therapeutic LNPs is similar to the LNPs investigated here, this would imply that the average copy number should be lower when using a larger cargo molecule. Therefore, considering that saRNA molecules typically consist of more than 10,000 nucleotides, saRNA-LNPs such as in a recently approved saRNA-based vaccine against SARS-CoV-2 are expected to be well suited for direct RNA counting using *in situ* staining as the saRNA size is much larger than the 20 nm gap for the fluid exchange.^45,46^ Furthermore, in the case of LNPs co-loaded with different pre-formulation labeled cargoes, the same direct counting approach is expected to be applicable. This would enable detailed characterization of LNPs containing complex cargoes, such as co-loaded Cas9-encoding mRNA and guide RNA in CRISPR gene editing systems. Thus, direct counting is expected to be applicable to a broad range of cargo sizes and types, with high accuracy expected in the regime of 5 or fewer cargo molecules.

Together, our results show that precise quantification of mRNA copy number is possible using single-molecule confinement for LNP lysis and labeling. We have validated our method using two distinct LNP formulations, but this approach could be applied to characterize LNP payload in the context of numerous formulation parameters, such as the ratio of lipid components, the choice of ionizable lipid(s), and the length of RNA cargo. The presented method for direct cargo counting is applicable to a broad range of clinically important LNP formulations and is complimentary to mRNA estimation based on rescaling the fluorescence intensity of the encapsulated RNA. This approach to mRNA counting can provide LNP developers with precise analytics of mRNA copy number and size at the single-particle level.

## Supporting Information

Supplementary Information is provided, with the following topics:

- Additional experimental details, methods, validation measurements and parameter optimization using computer simulations.
- Additional interpretation of the size-loading and Cy5-labeled mRNA-LNP results.

## Acknowledgment

Cryo-TEM grid preparation and data collection was performed at the High Resolution Macromolecular Electron Microscopy (HRMEM) facility at the University of British Columbia (https://cryoem.med.ubc.ca). We thank Claire Atkinson, Amy Wo, Barathy Deivanayaga, Liam Worrall and Natalie Strynadka. HRMEM is funded by the Canadian Foundation for Innovation and the British Columbia Knowledge Development Fund.

## TOC Graphic

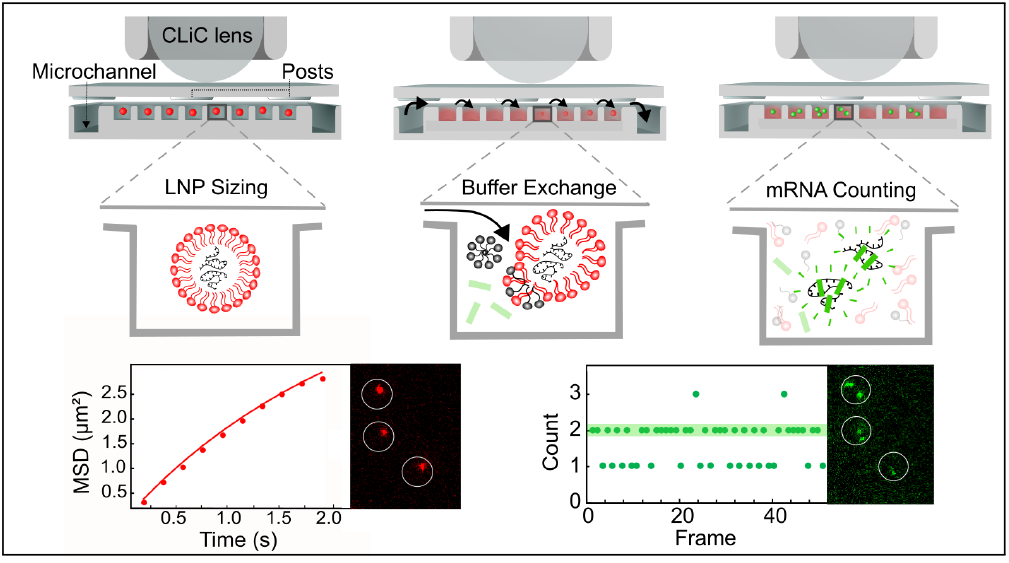

