## Supplementary Material for "Direct Counting of mRNA Copies Inside Individual Lipid Nanoparticles Using In Situ Lysis and Labeling"

### Contents

**S1 Experimental Protocol** **S3**

S1.1 CLiC Imaging flow-cell . . . . . . . . . . . . . . . . . . . . . . . . . . . . . S3

S1.2 Reagent optimization . . . . . . . . . . . . . . . . . . . . . . . . . . . . . . S4

S1.3 Acquisition . . . . . . . . . . . . . . . . . . . . . . . . . . . . . . . . . . . S5

**S2 Data analysis** **S6**

S2.1 Fluorescence denoising . . . . . . . . . . . . . . . . . . . . . . . . . . . . . S6

S2.2 Local maxima detection . . . . . . . . . . . . . . . . . . . . . . . . . . . . S8

S2.3 K-Max counting . . . . . . . . . . . . . . . . . . . . . . . . . . . . . . . . . S8

S2.4 2D Gaussian fitting . . . . . . . . . . . . . . . . . . . . . . . . . . . . . . . S11

S2.5 Mean square displacement analysis for particle sizing . . . . . . . . . . . . S11

S2.6 Power law analysis of particle size and loading . . . . . . . . . . . . . . . . . S12

**S3 mRNA-LNP Formulation** **S14**

S3.1 Components . . . . . . . . . . . . . . . . . . . . . . . . . . . . . . . . . . S14

S3.2 Preparation . . . . . . . . . . . . . . . . . . . . . . . . . . . . . . . . . . S14

S3.3 Characterization . . . . . . . . . . . . . . . . . . . . . . . . . . . . . . . . S15

**S4 RiboGreen Controls** **S16**

S4.1 Sensitivity to individual mRNA . . . . . . . . . . . . . . . . . . . . . . . . S16

S4.2 Specificity for mRNA . . . . . . . . . . . . . . . . . . . . . . . . . . . . . . S17

S4.3 Detectability relative to Cy5 . . . . . . . . . . . . . . . . . . . . . . . . . . S17

S4.4 Non-correlation of mRNA intensity and diffusivity . . . . . . . . . . . . . . . S19

**S5 Lysis experiments with Cy5-labeled mRNA** **S20**

S5.1 mRNA counting . . . . . . . . . . . . . . . . . . . . . . . . . . . . . . . . . S20

S5.2 Correlation of encapsulated Cy5 intensity and mRNA copy number . . . . . S23

**S6 Simulations of lysed mRNA** **S24**

S6.1 Simulated data generation . . . . . . . . . . . . . . . . . . . . . . . . . . S24

S6.2 Effect of SNR . . . . . . . . . . . . . . . . . . . . . . . . . . . . . . . . . . . . S25

S6.3 Effect of intensity adjustment . . . . . . . . . . . . . . . . . . . . . . . . S26

S6.4 Effect of noise background . . . . . . . . . . . . . . . . . . . . . . . . . . S29

S6.5 Effect of well size . . . . . . . . . . . . . . . . . . . . . . . . . . . . . . . . S30

S6.6 Effect of exposure time . . . . . . . . . . . . . . . . . . . . . . . . . . . . S32

S6.7 Effect of number of frames . . . . . . . . . . . . . . . . . . . . . . . . . . S32

**References** **S34**

### S1 Experimental Protocol

#### S1.1 CLiC Imaging flow-cell

The CLiC flow-cell is comprised of two 25 mm x 25 mm glass coverslips (Ted Pella, product no.260452) with thickness 200 ± 10 μm joined by a 30 μm thick double-sided adhesive (Nitto Denko, product no. 5603). The top coverslip contains an array of cylindrical posts (10 μm diameter, 20 nm thickness, 20 μm spacing) which form a nano-slit between the top and bottom coverslips for reagent exchange.[1] The bottom coverslip is patterned with an array of microwells (5 μm diameter, 500 nm depth, 3 μm spacing) which is connected to a micro-channel (20-30 μm depth, 200 μm width) to enable reagent exchange (Fig. 1 in the main text).

Prior to assembly, both coverslips underwent a passivation protocol as previously described[2,3] using polyethylene glycol (PEG) to minimize interactions between the LNPs and the glass surfaces. Briefly, coverslips were sonicated in acetone followed by isopropyl alcohol before being cleaned in a 3:1 (v/v) piranha solution of sulfuric acid ($H_2SO_4$, CAS: 7664-93-9, Thermo Fisher Scientific) and hydrogen peroxide ($H_2O_2$, CAS: 7722-84-1, Thermo Fisher

Scientific) for 45 min. The coverslips were then etched with KOH (1 M) for 10 min and dried with pressurized $N_2$ gas. A layer of 3-(aminopropyl)triethoxysilane (APTES, CAS: 919-30-2, Thermo Fisher Scientific) was then deposited onto the coverslips using a vacuum desiccator for 30 min. The coverslips were then heated on a hot plate for 20 min before adding 30 μL of N-hydroxylsuccinimide (NHS) functionalized 5 kDa polyethylene glycol valeric acid (PEG-SVA, Laysan Bio) in a 0.9 M solution of sodium sulphate ($Na_2SO_4$, CAS: 7757-82-6, Thermo Fisher Scientific). The coverslips were incubated with the PEG-SVA solution for 30 min before being rinsed with deionized water, dried, and assembled.

Once assembled, the CLiC flow-cell was fastened in a plastic chuck and placed below the curved surface of the CLiC pusher lens, which comprises a spherical lens held in a hollow cylinder. Using a piezoelectric motor, the CLiC lens was lowered to cause a downward deflection of the top coverslip until it came in contact with the bottom coverslip, creating a circular area of confinement ($r \approx 125$ μm) centered at the approximate center of the CLiC lens. Within this confinement area, each confined LNP and its mRNA cargo remained trapped in the same microwell for the duration of the reagent exchange across the nano-slit. To increase statistics, multiple 512×512 fields-of-views (FOVs) were imaged in the circular confinement region. For each experiment, the edges of the confinement region were empirically determined by manually inspecting the FOVs near the edge of the confined region for potential loss of confinement. This inspection was done for post-lysis mRNA videos as the size of the mRNA is smaller than that of the LNPs.

#### S1.2 Reagent optimization

The lysing agent (Triton® X-100) and RNA stain (Quant-iT™RiboGreen™RNA reagent) were tested at multiple concentrations to optimize the efficiency of the assay and quality of the imaging data. The concentration of Triton® X-100 detergent was optimized for completeness and rate of lysis. Five detergent concentrations (10%, 1%, 0.2%, 0.05%, and 0.01% (w/w)) were tested in the LNP lysis protocol depicted in Fig. 1. The 10%, 1%, and 0.2% detergent solutions produced similar rates and completeness of lysis, whereas the 0.05% solution produced a lesser degree of complete lysis and the 0.01% solution was incapable of lysing LNPs. Note that since the lysis occurred during fluid exchange, no further dilution of the lysis solution occurred, which made the concentration similar to that of other investigations.[4] Therefore, to minimize reagent usage and maximize completeness of lysis, a solution of 0.2% (w/w) Triton® X-100 was used as the lysis buffer for all subsequent experiments.

The concentration of Quant-iT™RiboGreen RNA reagent was optimized to maximize the intensity of labeled RNA while minimizing the fluorescence background due to dye interactions with the glass flow-cell. Four dilutions of RiboGreen (1:100, 1:200, 1:500, and 1:1000) were tested in the LNP lysis protocol depicted in Fig. 1. The dilutions of 1:100 and 1:200 produced poor signal-to-noise ratio (SNR) due to high background, whereas the 1:1000 also produced poor SNR due to insufficient dye molecules per mRNA. Therefore, a 1:500 dilution of RiboGreen for all subsequent experiments.

All reagent optimization and subsequent experiments were performed at room temperature.

#### S1.3 Acquisition

Image data was acquired using a Nikon Ti-E inverted microscope equipped with an Apo chromat TIRF 100× oil immersion objective lens (NA=1.49, WD=0.12 mm, FOV=22 mm) for wide-field imaging. The sample was excited by a co-aligned laser system comprising the Sapphire 488 nm LPX (100 mW) and OBIS 647 nm LX (120 mW). Emission wavelengths were separated by a long-pass dichroic filter (Chroma ET640lp) before being imaged on two separate Andor iXon Ultra 897 EMCCD cameras (16 $\mu$m/pixel, 512×512 pixels). A set of opaque slits (Thorlabs) was affixed in the beam path to truncate the beam area to the dimensions of one imaging FOV (22x22 mm) to prevent photobleaching of fluorophores in adjacent FOVs prior to data acquisition.

To further minimize photobleaching and photonicking, laser powers were set at 0.7-1.1 mW (measured at objective) and the number of frames was calibrated based on the intended measurable: 10 frames for pre-lysis Cy5 intensity measurements, 50 frames for mRNA counting, and 130 frames for LNP diffusivity measurements. Images were acquired with an exposure time of 20 ms to allow for high SNR with minimal motion blur. Excitation laser powers of 0.7-1.1 mW at the objective corresponds to 3-4 W/cm$^2$. This laser intensity, an exposure time of 20 ms, and the recording of 50 post-lysis frames was used to maximize SNR ($> 4$ per mRNA) while mitigating excessive photobleaching and potential photodamage of the fluorescently-labeled mRNA that can occur at high laser intensities.[5] Manual inspection of the data shows no noticeable occurrence of photodamage. The choice of acquisition parameters was assessed using simulations and was shown to enable accurate counting of individual fluorescently labeled molecules under the given experimental conditions (Section S6).

### S2 Data analysis

Prior to analysis, all imaging FOVs were manually aligned to a binary mask consisting of an array of circles to identify the centers of the microwells. These centers were then used to crop a 32×32 pixels region of interest around each well. Most microwells had a diameter of approximately 5 μm, but the precise well diameter also varied between different flow-cells due to fabrication differences. To account for this, the microwell diameter was measured for each experimental dataset in pixels and the binary well mask was generated with circles of the same diameter.

#### S2.1 Fluorescence denoising

After removing microwell videos outside the confined region (Section S1.1), a background subtraction was applied to all videos with a non-zero background, where the background for each image was calculated as a 32×32 array of the temporal median (e.g., median intensity over all frames) for each pixel. After background subtraction, the videos were further pre-processed to remove Poisson noise and residual background fluorescence using a U-Net-style convolutional neural network (Fig. S1). The U-Net was first published in the context of biomedical image processing,[6] but it is also applicable to microscopy data for tasks such as denoising and tracking.

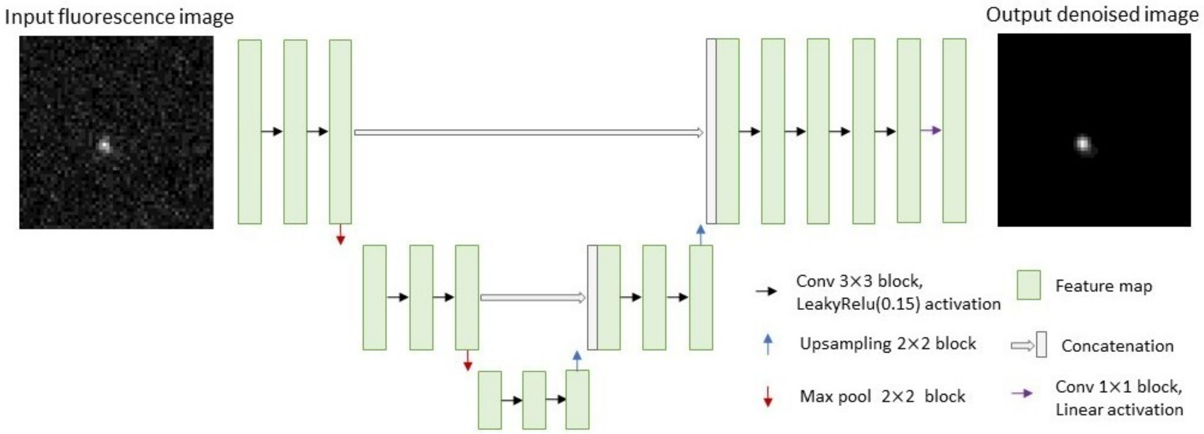


Figure S1: Architecture of U-Net for Denoising. The U-Net inputs a fluorescence image of size (32 × 32) pixels. The image is downsampled using convolutions and max pooling before being upsampled. Skip connections occur at each layer (e.g., row) to concatenate the downsampled and upsampled feature maps. The convolutional kernel size is 3 × 3.

The training data for the U-Net was generated using the DeepTrack 2.0 package.[7] Briefly, between 0-10 fluorescent point sources were simulated at a randomized intensity, planar position, and depth within a 32×32 pixels image. Poisson noise was added to the images using a randomly drawn $\lambda$ value from a range designed to mimic the distribution of SNRs observed in the experimental data. In addition to Poisson noise, a random linear background gradient was added to replicate the non-uniformity of staining across the well array seen in the experimental data and a weak ring-shaped signal was added to mimic sticking of the stain to the microwell edges. The numerical aperture and optical aberrations were also simulated to match the experimental data. To assess the performance of the denoising network, a subset of simulated training videos were manually inspected before and after denoising. The dynamic range of the network was further validated by generating confusion matrices to show the counting accuracy over various simulated mRNA counts at a range of SNRs (Section S6.2). The network was trained with Tensorflow 2.10.0 using mean absolute error and an Adam optimizer (learning rate=8e-5).[8]

#### S2.2 Local maxima detection

After denoising, particle detection was performed using the Crocker-Grier algorithm[9] and implemented using a modified version of *trackpy* in Python 3.12.[10] Briefly, a rough local-maxima search is performed by looking for pixels that are higher intensity than all neighboring particles. Then, a refinement of the pixel weight is applied according to:

$$\vec{\boldsymbol{\epsilon}} = \frac{1}{m} \sum_{i,j \leq w^2} \vec{\mathbf{x}}_{ij} A[x+i, y+j], \tag{S1}$$

where $\vec{\mathbf{x}}_{ij}$ is defined as the vector with array index $i, j$ in the image, $\vec{\boldsymbol{\epsilon}}$ is defined as the refinement of a particle, and $w$ is defined as the rough area of the particle. Lastly, $m$ is defined as the discrete integral over the image, $A[x, y]$ in the region around the particle.

In addition to providing data for post-lysis mRNA counting, the local maxima detections were used to monitor the completion of lysis in the well. Specifically, the detected signal in the lipophilic dye channel was compared pre- and post-lysis and only wells in which the post-lysis signal was less than 20% of the initial signal were retained for the counting analysis.

#### S2.3 K-Max counting

After identifying the local intensity maxima, a counting algorithm was applied to 1) identify microwells with a single LNP and 2) estimate the number of mRNA molecules in each microwell. Once denoised, wells with a single LNP were easily identified by taking the median number of detections over all frames, however, a custom estimator was needed for accurate determination of the number of mRNA per microwell. This is because the number of mRNA detected in each frame can vary as a result of both Poisson statistics and the potential overlap between mRNA molecules in the 2D projection of the well (Fig. S2). To accurately estimate the number of particles in a confined volume across multiple frames, a robust metric is therefore required.

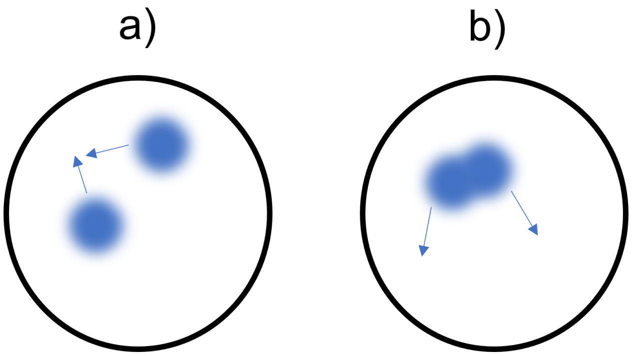


Figure S2: Multiple diffusing mRNA in the microwell may be adequately separated (a) or overlapping (b) in different imaging frames.

Simulations of mRNA molecules (Section S6) in microwells (5 μm diameter) show that traditional estimators such as mean or median are likely to underestimate the true number of mRNA in a microwell (Fig. S3, top left-middle). To mitigate this, we propose using an adapted maximum as an estimator. Simulations of particles with an SNR of 6 show that in the moderate copy number regime (e.g., 4-5), a traditional maximum is more accurate than the mean or median (Fig. S3, top right). However, in the low copy number regime (e.g., 0-3), the maximum tends to overestimate the count due to noise-driven outliers. To balance robustness with sensitivity, we introduce and define the **K-Max** estimator. Let $W$ be a list of detections of some arbitrary length. After sorting $W$ in descending order (e.g., largest to smallest), we can take $X_W$ to be a sublist of $W$. The length of $X_W$ can be defined by $k$, which will determine the number of ordered detections to consider. Taking $X_W^{[:k]}$ as the sublist of the top $k$ counts, K-Max is defined as the median of $X_W^{[:k]}$. For example, if $X_W = [5, 5, 4, 3, 2]$ and $k = 3$, then $X_W^{[:3]} = [5, 5, 4]$ and K-Max would be 5.

The K-Max estimator offers a tunable compromise between the stability of the median and the robustness of the maximum. Simulations show that the K-Max estimator significantly improves the counting accuracy in the low to moderate copy number regime for molecules with an SNR of 6 (Fig. S3, bottom). For high copy number estimation, the K-Max estimator was much less effective; therefore, counts in this regime were rescaled based on the intensity over the entire microwell for improved accuracy (Section S6.3). The K-Max statistic was tested for $k$ values of 2, 3, and 4 and a $k$ of 3 showed the best overall accuracy (Fig. S3, bottom). To avoid overcounting in the case of empty wells or single mRNA without overlap, the K-Max statistic was only used when the mean count was >1. When the mean count was <1, the overall median was used as an effective statistic.

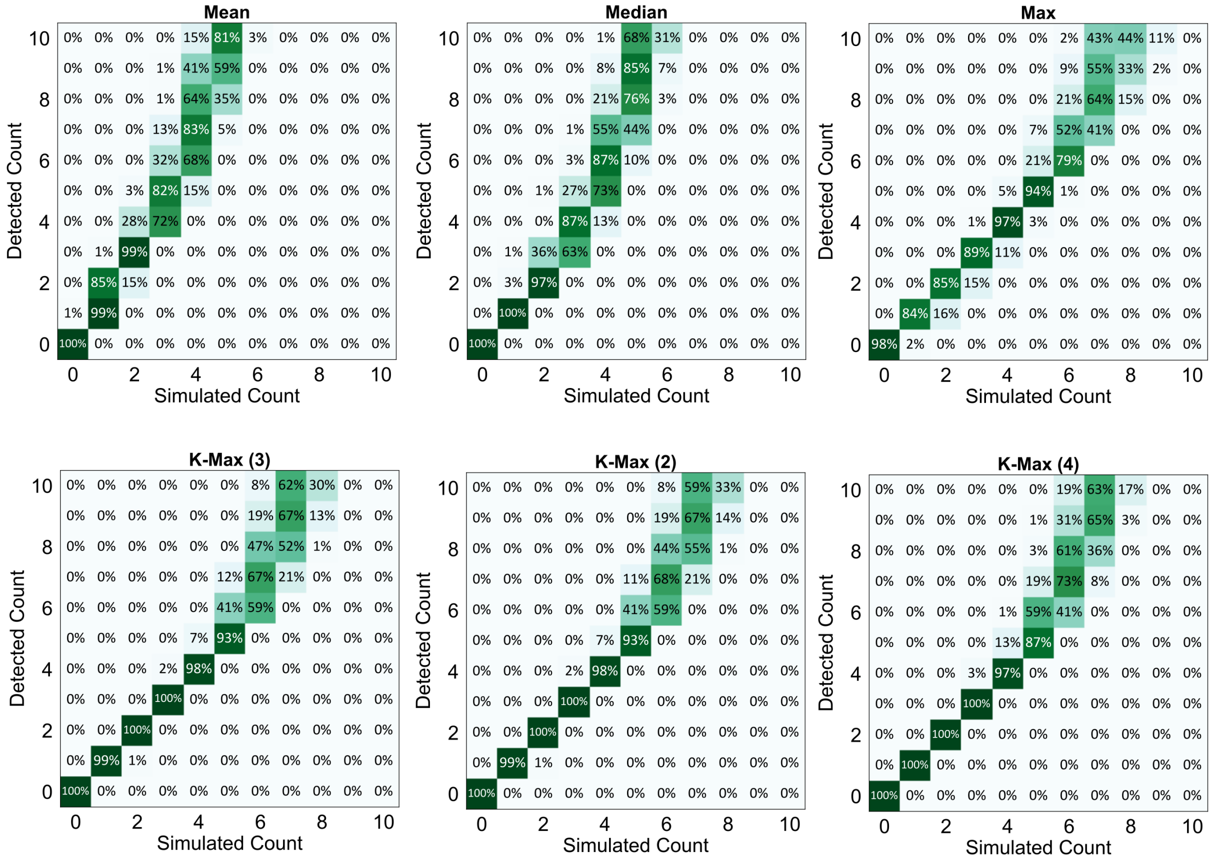


Figure S3: Confusion matrices showing the accuracy of automated counting on simulated mRNA videos using the traditional metrics of mean, median, and maximum (top) versus using the customized K-Max statistic (bottom) with $k$ values of 2, 3, and 4.

For each well, a check was also performed to detect the presence of any static (e.g., non-diffusing) mRNA signals that were removed during the background subtraction step.

Non-diffusing mRNA signals were detected by first applying the local maxima detection to the temporal intensity median (defined in Section S2.1), then filtering the detections using an empirically deduced intensity threshold to prevent overcounting.

#### S2.4 2D Gaussian fitting

The intensity signals of free RiboGreen-labeled mRNA and encapsulated Cy5-labeled mRNA were quantified by fitting a symmetric 2D Gaussian, as given by:

$$f(x, y) = A \cdot e^{-\frac{(x-x_0)^2+(y-y_0)^2}{2\sigma^2}}, \tag{S2}$$

using least squares.

Prior to applying a Gaussian fit, the pre-lysis Cy5 videos underwent additional pre-processing. Using the described algorithm for local maxima detection (Section S2.2), the centroid of the signal in each frame was located. A 12×12 pixel area was cropped around the centroid in each frame and the 10 cropped images were averaged to artificially improve the SNR.[11]

#### S2.5 Mean square displacement analysis for particle sizing

Denoised well videos with a median count of one LNP were retained for analysis. These wells were recounted and tracked with a constraint on the number of detections per frame to be one, which was used to generate the final 130-frame particle trajectory.

From this trajectory, the mean square displacement (MSD) was calculated over 10 consecutive positions and fit to the model of 2D confined diffusivity described by Bickel et al.[12] as applied in prior publications using CLiC.[3,13,14]

All MSD fits were assessed for goodness of fit and only those adhering to the following criteria were retained for sizing analysis: a) local tracking uncertainty <0.5 µm$^2$, b) estimated well radius $r$ in the range of 0.5 µm $< r <$ 15 µm, and c) estimated diffusivity $d$ in the range of 0.1 μm$^2$/s $< D <$ 80 μm$^2$/s.

A diffusion coefficient was then estimated from each retained MSD fit and used to estimate particle size using a modified Stokes-Einstein equation to correct for confinement effects. When particles are proximal to a surface, they experience a drag force that decreases the apparent diffusivity.[13] We correct for this effect in our parallel-plane geometry by including a correction factor $\lambda$ to the Stokes-Einstein relation as in:

$$D_{||} = \frac{\kappa_B T}{6\pi\eta a\lambda} = \frac{D_0}{\lambda}, \tag{S3}$$

where $D_{||}$ is the proximity-corrected diffusivity, $\kappa_B$ is the Boltzmann constant, $T$ is the temperature, $\eta$ is the viscosity, and $a$ is the hydrodynamic radius of the LNP.

A $\lambda$ value is calculated for a given radius estimate using Eq.S4, where $z$ is half the depth of the cylindrical confinement volume (e.g., distance from the center of the particle to the parallel surfaces). For the materials used in this study, $z = 500$ nm in all experiments. The raw $\lambda$ value is calculated for a given size using:

$$\lambda = 1 - 1.004\left(\frac{a}{z}\right) + 0.418\left(\frac{a}{z}\right)^3 + 0.210\left(\frac{a}{z}\right)^4 - 0.169\left(\frac{a}{z}\right)^5, \tag{S4}$$

which is then compared to a predefined set of possible correction values ranging from 0.05 to 1.00 (step size=0.05) and the closest $\lambda$ from the set to the raw estimate is used as the final $\lambda$ value.

#### S2.6 Power law analysis of particle size and loading

In addition to the Pearson correlation analysis reported in the main text, the relationship between particle size and loading was investigated using power law analysis. First, the particle size data was split into six bins of equal size, where the bin size was chosen to ensure approximately equal data points per interval in terms of hydrodynamic radius. The mean particle size and the corresponding mean mRNA copy number were then calculated for each bin and plotted on a set of log-log axes. As shown in Fig. S4, this produces a distribution that can be fit with a linear relationship, yielding a slope of 0.21 $\pm$ 0.03 for LNPs formulated in NaOAc buffer and a slope of 0.35 $\pm$ 0.03 for LNPs formulated in NaCit buffer. This indicates that the relationship between particle size and mRNA loading in both formulations may follow a power law, which can be expressed as $\mu = \beta R^{\alpha}$, where R is the hydrodynamic radius of the LNP.

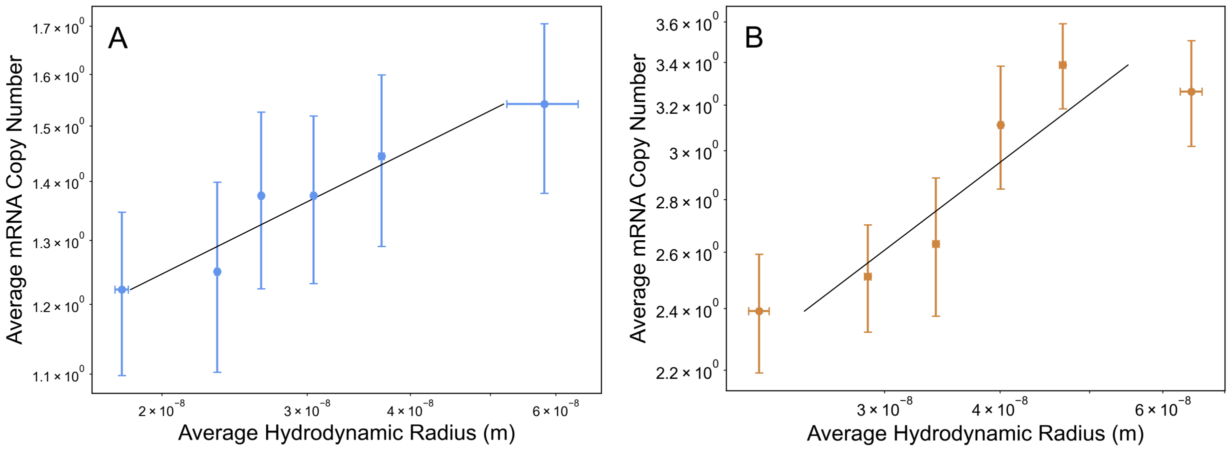


Figure S4: Log-log plots of average mRNA copy number as a function of average particle radius for LNPs formulated in (A) NaOAc (n=415) and (B) NaCit (n=270) buffer. Error bars represent the standard deviation of the mean. The slopes of the linear fits are 0.21 $\pm$ 0.03 and 0.35 $\pm$ 0.03 for NaOAc and NaCit, respectively.

For both LNP formulations containing unlabeled mRNA cargo, the distributions of mRNA copy number (Fig. 2B in the main text) were reminiscent of a Poisson distribution with low variance (e.g. $\lambda < 2$). A resemblance to a low-variance Poisson distribution was also observed in the mRNA copy number data for LNP formulations containing Cy5-labeled mRNA, as reported in Section S5. Therefore, setting $\mu$ as the mean of a hypothetical Poisson distribution, we can estimate the values of $\beta$ and $\alpha$ to describe mRNA loading into LNPs as a stochastic Poisson-based process.

We used a model of maximum likelihood to estimate the following values: $\alpha_{\mathrm{NaOAc}} = 0.278$, $\beta_{\mathrm{NaOAc}} = 169$, $\alpha_{\mathrm{NaCit}} = 0.331$, and $\beta_{\mathrm{NaCit}} = 820$. The values of $\alpha$ are in strong agreement with the graphs in Fig. S4 and the increased value of $\beta$ for the NaCit dataset further supports the increased loading of mRNA in LNPs formulated using NaCit than those formulated using NaOAc. This is tentative evidence that mRNA loading into LNPs may be modeled as a Poisson process, however, longer particle track lengths and a larger dataset of single-particle size and mRNA copy number measurements is needed to support these preliminary findings. For example, statistical uncertainty when estimating the hydrodynamic radius, which here is present due to the finite track length, could potentially underestimate $\alpha$. Therefore, more detailed measurements and comprehensive modeling of mRNA-LNP formation at the molecular level could aid in elucidating the biophysical basis of this process.

### S3 mRNA-LNP Formulation

#### S3.1 Components

The lipids 1,2-distearoyl-sn-glycero-3-phosphorylcholine (DSPC) and 1,2-dimyristoyl-rac-glycero-3-methoxypolyethylene glycol-2000 (PEG-DMG) were purchased from Avanti Polar Lipids. The ionizable lipid DLin-MC3-DMA (MC3) was synthesized in Dr. Marco Ciufolini's lab at the University of British Columbia. Lipophilic dyes 1,1-Dioctadecyltetramethylindodicarbocyanine (DiD) and 3,3'-Dioctadecyloxacarbocyanine perchlorate (DiO) were purchased from ThermoFisher Scientific. Unlabeled mRNA (m1Ψ-modified, catalog No. R1004) and Cy5-labeled mRNA (5-moUTP-modified, catalog No. R1010) molecules encoding Firefly luciferase (1921 nt) were purchased from APExBio. Triton® X-100 was purchased from MP Biomedicals LLC and Quant-iT™RiboGreen RNA reagent was purchased from Thermo Fisher Scientific.

#### S3.2 Preparation

The lipid components (MC3/Chol/DSPC/PEG-DMG/DiX) were dissolved in ethanol at a molar ratio of 50/38.5/10/1.5/1 to a final lipid concentration of 10 mM, where DiX was either DiD or DiO. DiD was added to formulations containing unlabeled mRNA and DiO was added to formulations containing Cy5-labeled mRNA to ensure spectral separation between the lipid and mRNA channels. The LNPs were then formed by mixing the lipid components with an aqueous buffer (pH 4) containing the mRNA via a T-junction mixer[15,16] (3:1 aqueous:organic phase (v/v) at 20 mL/min flow rate) and then dialyzed overnight against 1X PBS (pH 7.4). The aqueous buffer was chosen as 25 mM NaOAc or 300 mM NaCit. The mRNA was added at a concentration to achieve a nitrogen-to-phosphate (N/P) ratio of 6. Prior to imaging, lipid nanoparticles were diluted in 1X PBS (pH 7.4) to an empirically determined concentration to confine a single particle in the majority of wells, as previously described.[1,13] Two sample replicates were prepared for each formulation and each sample replicate was analyzed a minimum of three times within 8 days of the formulation date, in accordance with stability measurements from prior studies.[17–20].

#### S3.3 Characterization

Prior to the CLiC measurements, LNP size measurements were taken for at least one sample replicate via dynamic light scattering (DLS) using a Malvern Zetasizer Nano ZS and average sizes were reported as intensity-weighted averages (Table S1). The discrepancy between the size measurements from DLS and CLiC can be explained by the type of average that each method reports. In the DLS data, we report an intensity-weighted average, which favours larger particle sizes due to their increased scattering, whereas our CLiC-based measurements report a number average, which weighs all particle sizes equally. In Table S1, note that only the LNPs with post-labeled mRNA are shown in the main text. Cryogenic transmission electron microscopy (cryoTEM) was also used to provide LNP structure information. LNPs were prepared for cryoTEM as previously described.[3] Briefly, LNPs were concentrated to 15−20 mg/mL total lipid before being plunge frozen on copper grids using a FEI Mark IV Vitrobot. Grids were imaged on a FEI LaB6 G2 TEM by the High Resolution Macromolecular Cryo-Electron Microscopy facility at the University of British Columbia (UBC) under 55,000X magnification. Finally, the RNA encapsulation efficiency was measured using the Quant-it® RiboGreen Reagent and RNA Assay Kit (Thermofisher) for the LNP formulations containing unlabeled mRNA. For both the NaOAc and NaCit samples, the encapsulation efficiency was >99%, which is in agreement with typical values for Onpattro®-based LNP formulations.

Table S1: Size characterization of mRNA-LNP formulations. For the DLS diameter, the intensity average is used.

| Formulation Buffer | Lipid Dye | mRNA Dye | DLS | | CLiC |
|---|---|---|---|---|---|
| | | | PDI | Diameter (nm) | Diameter (nm) |
| NaOAc | DiO | Cy5 | 0.144 | 65.7 ± 1.5 | n/a |
| NaOAc | DiD | RiboGreen (post-labeled) | 0.184 | 65.3 ± 3.5 | 60.1 ± 1.0 |
| NaCit | DiO | Cy5 | 0.195 | 161.4 ± 0.4 | n/a |
| NaCit | DiD | RiboGreen (post-labeled) | 0.083 | 78.0 ± 0.3 | 78.4 ± 1.8 |

### S4 RiboGreen Controls

The suitability of RiboGreen for in situ mRNA labeling was assessed in terms of its sensitivity and specificity for mRNA as well as the detectability of RiboGreen-labeled mRNA relative to Cy5-labeled mRNA. Furthermore, the assumption that individual intensity peaks correspond to individual mRNA was validated by measurements of free RiboGreen-labeled mRNA showing that intensity does not correlate with diffusivity.

#### S4.1 Sensitivity to individual mRNA

To assess the sensitivity of RiboGreen (e.g., its ability to detect single mRNA molecules), RNA detections based on RiboGreen labeling and detection based on covalent Cy5 labeling were compared. LNPs loaded with Cy5-labelled mRNA were isolated and imaged under CLiC to quantify the proportion of empty and non-empty particles based on the detection of encapsulated Cy5 signals. Particles were then lysed and stained with RiboGreen as in the typical protocol. Using manual inspection, wells were classified as empty or non-empty (e.g., containing mRNA) using the Cy5 and RiboGreen channels separately and the correspondence between detections was plotted as a confusion matrix. Per Fig. S5A, 98% of wells with no Cy5 detection also had no detection of RiboGreen and 91% of wells with a Cy5 detection also had a RiboGreen detection. The 9% non-agreement between Cy5 and RiboGreen may be attributed to the non-uniformity of staining across the well array, where wells in the center of the confined region were less permeated by the stain, or incomplete LNP lysis, resulting in lower intensity RiboGreen labeling that was potentially not detectable by manual inspection. This could come from the tradeoff between nano-slit size and microfluidic permeation, where having a small nano-slit with surface PEGylation prevents mRNA escape, but also hinders RiboGreen and Triton X-100 from entering the wells. This suggests that the fraction of empty LNPs might be slightly overestimated, while the mRNA number distribution of the detected loaded LNPs should remain accurate.

#### S4.2 Specificity for mRNA

The specificity of RiboGreen was validated by lysing and staining unloaded LNPs (e.g., no mRNA cargo) as in Fig. S5B. Using this method, 100% of wells (n=169) were correctly classified by the automated counting as containing zero mRNA, which shows that RiboGreen staining is specific to mRNA and unlikely to produce false detections due to interactions with the glass.

#### S4.3 Detectability relative to Cy5

In preliminary lysis experiments, individual mRNA both with and without Cy5 labels were imaged. The SNR of a single mRNA in the Cy5 and RiboGreen channels was quantified as an average over the first 10 frames and plotted as a histogram in Fig. S5C, which demonstrates that RiboGreen-labeled mRNA has a higher average SNR (8.6) than Cy5-labeled mRNA (4.7). The SNR of individual mRNA as a function of exposure time was also plotted in Fig. S5D to show the relative photobleaching rates of RiboGreen and Cy5. Paired comparisons of Cy5- and RiboGreen-labeled mRNA show that while initial SNR is comparable between these

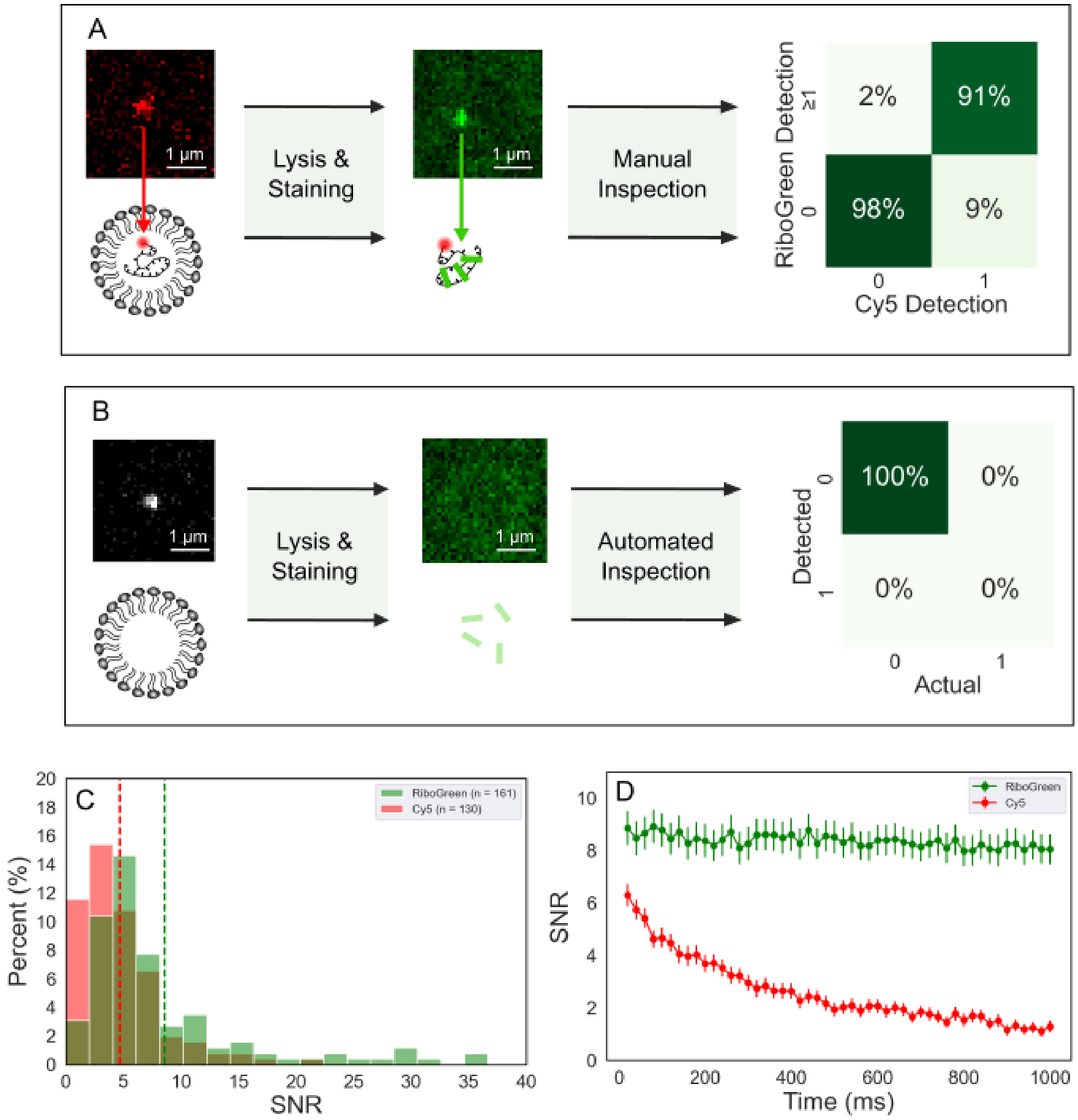


Figure S5: (A) Imaging schematic of Cy5-labeled mRNA cargo inside LNPs pre-lysis and RiboGreen-labeled mRNA post-lysis (left). Confusion matrix showing agreement between pre-lysis Cy5 and post-lysis RiboGreen detections with 98% agreement for empty wells ($n_{empty}$=697) and 91% for non-empty wells ($n_{non-empty}$=247) (right). (B) Experimental workflow of negative control for RiboGreen specificity using unloaded LNPs (left). Confusion matrix showing 100% accuracy of automated detection to classify empty wells ($n_{empty}$=169) (right). (C) Histogram of mean SNRs for RiboGreen- and Cy5-labeled mRNA over 10 frames of 20 ms exposure, measured as Gaussian-fit peak intensity over the standard deviation in the noise background. (D) SNR decay curves for RiboGreen- and Cy5-labeled mRNA during 1000 ms exposure ($n_{RiboGreen}$=161, $n_{Cy5}$=130).

labels, Cy5 bleaches significantly over the course of 1000 ms exposure whereas fluorescence intensity from RiboGreen remains fairly stable. This may be owed to stabilizing effects of Triton X-100 on RiboGreen intensity, as reported by Bizmark et al.[21]

#### S4.4 Non-correlation of mRNA intensity and diffusivity

Control measurements of free RiboGreen-labeled mRNA were performed to verify that RiboGreen intensity is uncorrelated with mRNA diffusivity. A 0.75 μL aliquot of Firefly Luciferase mRNA was defrosted on ice for 10 minutes then diluted 1:100 with 1X TE buffer. Then 1 μL of mRNA was added to a 1:1000 dilution of RiboGreen in 0.2% (w/w) Triton X-100 and incubated for 5 minutes to stain the mRNA. The RiboGreen dilution was optimized to maximize SNR as a function of mRNA staining and background fluorescence due to dye sticking on the glass. Stained mRNA was then flowed into the chuck and confined to be imaged for diffusivity measurements (130 frames, 20 ms exposure). The laser power was optimized to maximize SNR while mitigating photobleaching and photonicking of the RNA. The intensity of all single mRNA (n=578) was plotted against the RNAs' diffusivity to identify if the variation in RiboGreen intensity was due to aggregation of multiple mRNA or intrinsic variation in dye intercalation. Intensity was defined as the amplitude of a 2D Gaussian as fit to the mRNA intensity peak. Wells containing single diffusing mRNA with sufficient SNR for particle tracking were selected using the following criteria: a) K-Max count of 1, b) variance in counting <0.5 (to prevent the inclusion of noisy empty wells or wells with more than one mRNA), c) fitted well radius $r$ within 1 μm $< r <$ 15 μm, and d) local tracking error of <1 μm$^2$ per the MSD fit.

There is little correlation between single mRNA diffusivity and intensity (Fig. S6) indicating that the variation in labeling is due to the stochastic nature of RiboGreen intercalation and not RNA aggregation. The diffusivity distribution of the mRNA may be considered lower on average (median=10.5 μm$^2$/s) than expected for this sequence length (1921 nt). This may be the result of surface interactions between the mRNA and the flow-cell. The flow-cell is

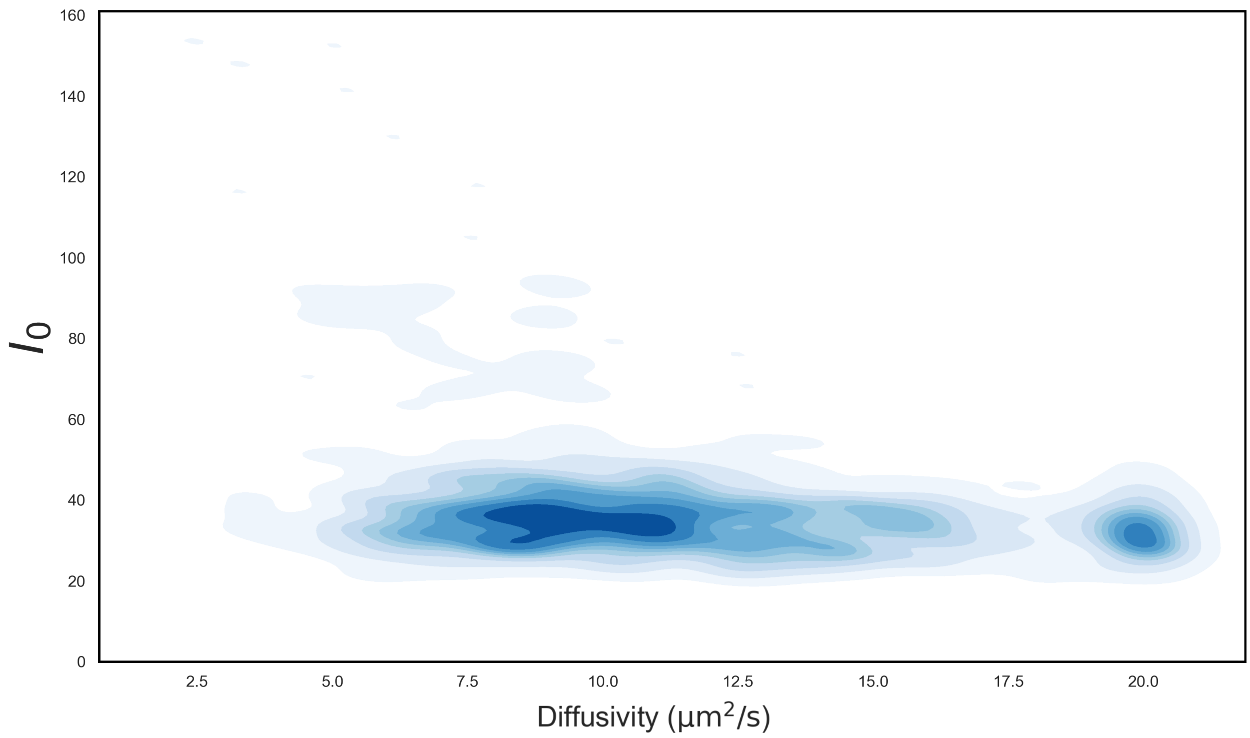


Figure S6: Density plot of individual mRNA intensities quantified via 2D Gaussian fit as a function of individual diffusivities (n=578).

intended to have a slight positive charge following passivation to reduce interactions with the LNPs, however, this charge may induce a drag force on the mRNA due to electromagnetic attraction. Another explanation could be that lysed lipids from the LNP in the post-lysis buffer bind to the mRNA and create a lipid-RNA complex with slower diffusion. Although it is unclear why the measured mRNA diffusivity is lower than expected, mRNA diffusion is not a primary focus of this work and the validity of individual mRNA counting is not challenged by this finding.

### S5 Lysis experiments with Cy5-labeled mRNA

#### S5.1 mRNA counting

While covalently attached labels such as cyanine dyes are useful for specific localization of RNA, the covalent attachment of fluorophores to RNA can reduce its negative charge and affects the size of the molecule.[22] Given the importance of RNA interactions with ionizable lipids during particle assembly and endosomal escape,[23] it is reasonable to assume that labeled RNA may behave differently than unlabeled RNA in these processes. Endosomal escape in particular is not well understood and considered a limiting step in the performance of RNA-LNPs.[24,25] Furthermore, the use of labeled RNA has also been linked to differences in the efficiency of transfection and translation.[26,27] Therefore, we compared the distribution of mRNA loading into LNPs using both unlabeled and Cy5-labeled cargo to investigate possible impacts on both the average copy number and spread in loading.

Fig. S7 shows the distribution of copy number using Cy5-labeled mRNA, which shares a similar shape (i.e., Poisson-like) to the distribution of copy number per the detection of RiboGreen-labeled mRNA. The average mRNA copy number ($n$) as measured using Cy5-labeled mRNA is similar to the average measured using RiboGreen-labeled mRNA in both the NaOAc ($n_{\mathrm{RiboGreen}} = 1.3$, $n_{\mathrm{Cy5}} = 1.8$) and NaCit ($n_{\mathrm{RiboGreen}} = 2.7$, $n_{\mathrm{Cy5}} = 2.6$) formulations. The primary difference between LNPs with Cy5-labeled and unlabeled cargo is the fraction of empty particles, where the unloaded fraction in the NaOAc formulation decreased from 35% to 25% when Cy5-labeled mRNA was used (Fig. S7B). Conversely, the unloaded fraction in the NaCit formulation increased from 20% to 31% (Fig. S7B). Thus, in all cases, the fraction of empty particles is around 20-35%, which is in line with previously reported values for LNPs with a similar size and cargo.[3]

We validated the method of lysing and counting Cy5-labeled mRNA to measure the unloaded fraction by taking measurements of Cy5 pre-lysis, as in Kamanzi et al.[3] Specifically, the unloaded fraction was measured using both the encapsulated Cy5 signal and the post-lysis diffusing Cy5-labeled mRNA signal. The unloaded fraction using both detection methods was within one standard error of the mean per the experimental replicates (Fig. S7B-C) showing the validity of the lysis counting method. Similar to the formulations with unlabeled mRNA, the Cy5-labeled mRNA-LNPs prepared in NaCit had a more polydisperse loading distribution due to a large fraction of highly loaded particles (e.g., >11% particles contained

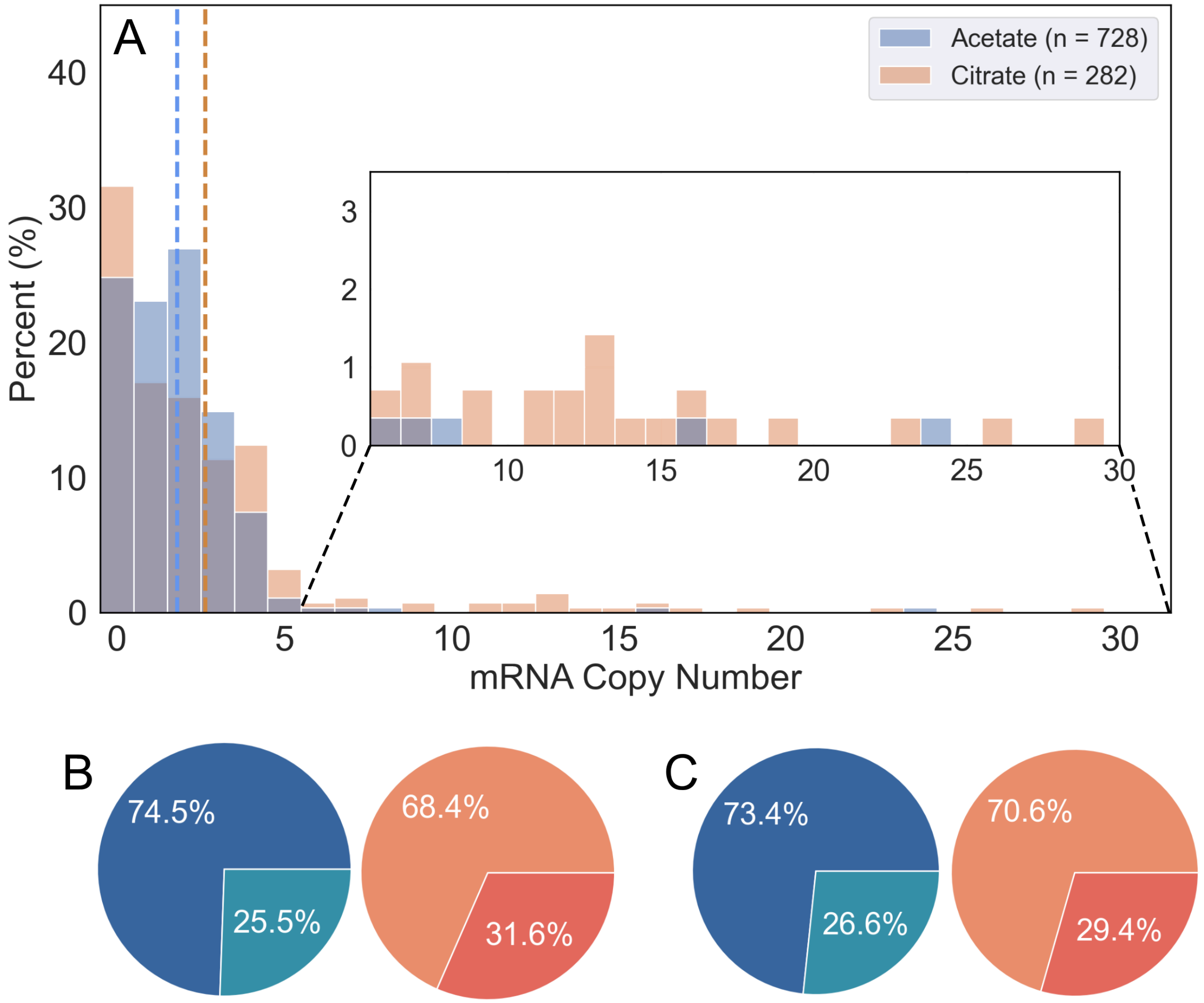


Figure S7: (A) Histogram of mRNA copy number per counting of Cy5-labeled mRNA for LNPs formulated in NaOAc (blue, n=728) and NaCit (orange, n=282) buffer. Piecharts of loading fraction for NaAOc (blue) and NaCit (orange) formulations based on detection of (B) post-lysis Cy5-labeled mRNA and (C) pre-lysis encapsulated Cy5-labeled mRNA.

>4 mRNA), as compared to the NaOAc formulation (e.g., <3% particles contained >4 mRNA). Overall, the distributions of mRNA copy number in LNPs with Cy5-labeled cargo are similar to those with unlabeled cargo, but differences in the percentage of unloaded particles and particles with high copy number may indicate that Cy5 labels influence mRNA encapsulation into LNPs.

#### S5.2 Correlation of encapsulated Cy5 intensity and mRNA copy number

The number of mRNA per LNP is commonly estimated by rescaling the measured intensity using the average intensity of a single labeled mRNA[3,24,28] Here we compare this number estimate with direct counting, which is measured pre-lysis, to the direct measurement of the number of mRNA determined post lysis. This allows us to investigate whether factors such as quenching and FRET impact mRNA number estimates when using intensity rescaling.

When comparing the intensity of the encapsulated Cy5 tags pre-lysis to the number of mRNA detected post-lysis, there is a proportional relationship as seen in Fig. S8. The relationship is not strictly linear, however, this is unsurprising given the limited ability of this method to quantify high copy number. On average, there is minimal quenching of Cy5 within the LNP prior to lysis, with <3% of LNPs having a non-detectable Cy5 signal pre-lysis that results in detectable mRNA post-lysis using Cy5. Therefore, the approach of using the encapsulated fluorophore intensity to estimate the mRNA copy number was relatively accurate in this study. This approach must still be assessed in the context of the specific LNP formulation at hand, where the extent of FRET can vary significantly as a function of the particle structure.[3]

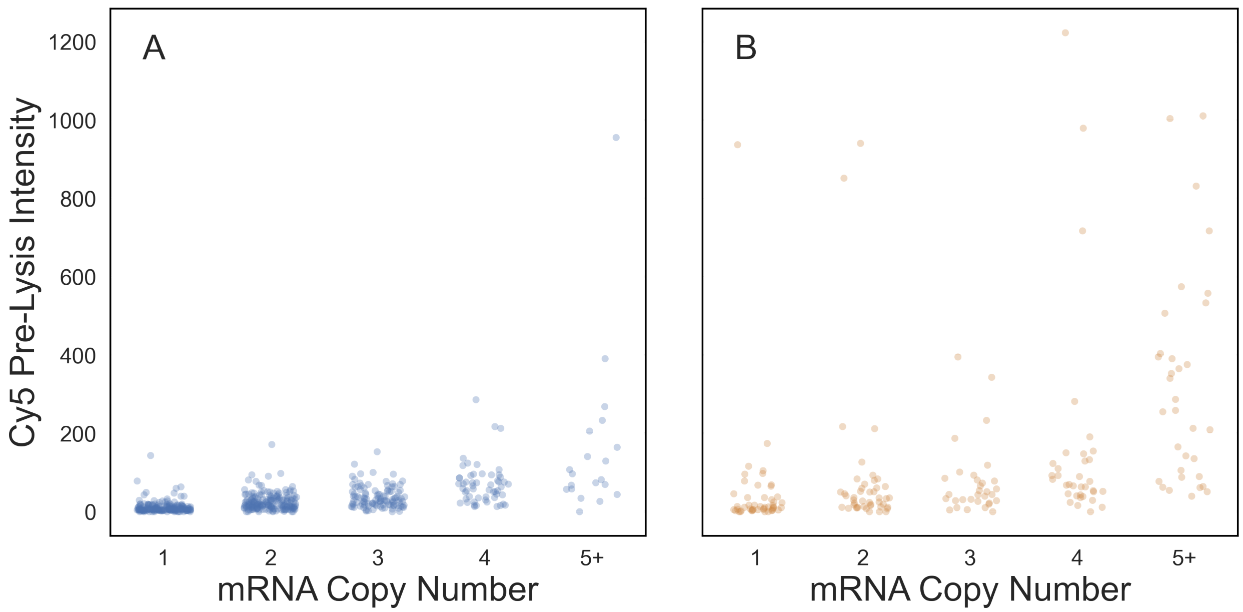


Figure S8: Scatter plots of pre-lysis Cy5 intensity signals as a function of mRNA copy number for LNPs formulated in (A) NaOAc buffer (n=728) and (B) NaCit buffer (n=282).

### S6 Simulations of lysed mRNA

Simulations were used to understand the strengths and limitations of both the experimental method and the automated mRNA counting software.

#### S6.1 Simulated data generation

Simulated particle videos were generated using the DeepTrack package designed for simulation and processing of microscopy data.[7] To obtain suitable intensity values for simulating mRNA, well videos were manually inspected and those containing a single mRNA were selected ($n_{\text{RiboGreen}}$=35, $n_{\text{Cy5}}$=38). For each video, a 2D Gaussian was fit to the peak in each frame and the the maximum amplitude in all frames was averaged to obtain a single intensity value. A 1D Gaussian distribution was then fit to the distribution of peak intensity values from all videos. Simulated intensity values were then drawn from a uniform probability distribution over the 65% confidence interval from the fitted 1D normal distribution. Using this intensity distribution, all videos could be generated with a specified SNR by choosing a random intensity and adding the appropriate level of Poisson noise to produce the expected SNR. In this case, SNR was defined as $\frac{I_0}{\sigma}$ where $I_0$ was the peak intensity of the particle modeled as a 2D Gaussian and $\sigma$ was the standard deviation in the noise background. The photobleaching of RiboGreen and Cy5 labels was also incorporated into the simulations to understand the effect of imaging time. Well videos containing one or more mRNA were identified using the automated counting software and denoised ($n_{\mathrm{RiboGreen}}$=43, $n_{\mathrm{Cy5}}$=65). For each denoised video, the sum of all pixels in the well area was measured for each frame and an exponential model of the form $I = I_0 e^{-kt}$ was fit to the time-series of summed intensities, where $I_0$ is the initial amplitude and $k$ is the decay constant. The same fitting process was applied to the pixel data outside the well area (e.g., background) to monitor changes in noise such that an overall decay constant for SNR could be ascertained. All fits were manually inspected and those showing a high goodness of fit were used to assess the rates of photobleaching ($n_{\mathrm{RiboGreen}}$=11, $n_{\mathrm{Cy5}}$=43). The average decay $k$ of the SNR of the RiboGreen-labeled mRNA was 0.15$\mathrm{s}^{-1}$ and for the Cy5-labeled mRNA it was 0.39 $\mathrm{s}^{-1}$. A conservative $k$ value of 0.39 $\mathrm{s}^{-1}$ was therefore used as the SNR decay constant for all simulations to maximize the number of analyzable wells in response to possible photobleaching.

#### S6.2 Effect of SNR

There is variable SNR among individual RiboGreen-mRNA due to 1) differences in secondary structure causing different levels of RiboGreen intercalation, 2) non-uniform spatial distribution of RiboGreen across the well array and across experimental replicates. Therefore, we investigated the accuracy of the counting software as a function of this key variable. For SNRs of 8, 6, and 4, the analysis software was highly accurate (>98%) in classifying between 0 to 4 mRNA per well using K-Max with $k$=3 (second column of Fig. S9). Above 4 mRNA per well, the accuracy of the counting decreases to between 90-93% for all three SNRs due to positional overlap of mRNA intensity peaks in the confined volume. In the experimental data, SNRs <4 comprised <5% and <3%, respectively, of the RiboGreen-labeled mRNA signals as reported in the main figures, thus high accuracy in counting can be expected for the majority of the data. For low-SNR data (e.g., <4), the maximum performs better as a counting metric than K-Max for 0-3 counts (bottom row of Fig. S9). Given that 51% of single Cy5-labeled mRNA data had an SNR <4 (Fig. S5B), the maximum was used in place of K-Max as a counting metric for Cy5-labeled mRNA data for more accurate counting. In general, imaging data with SNR <2 is prone to undercounting, which highlights the importance of having a good imaging SNR.

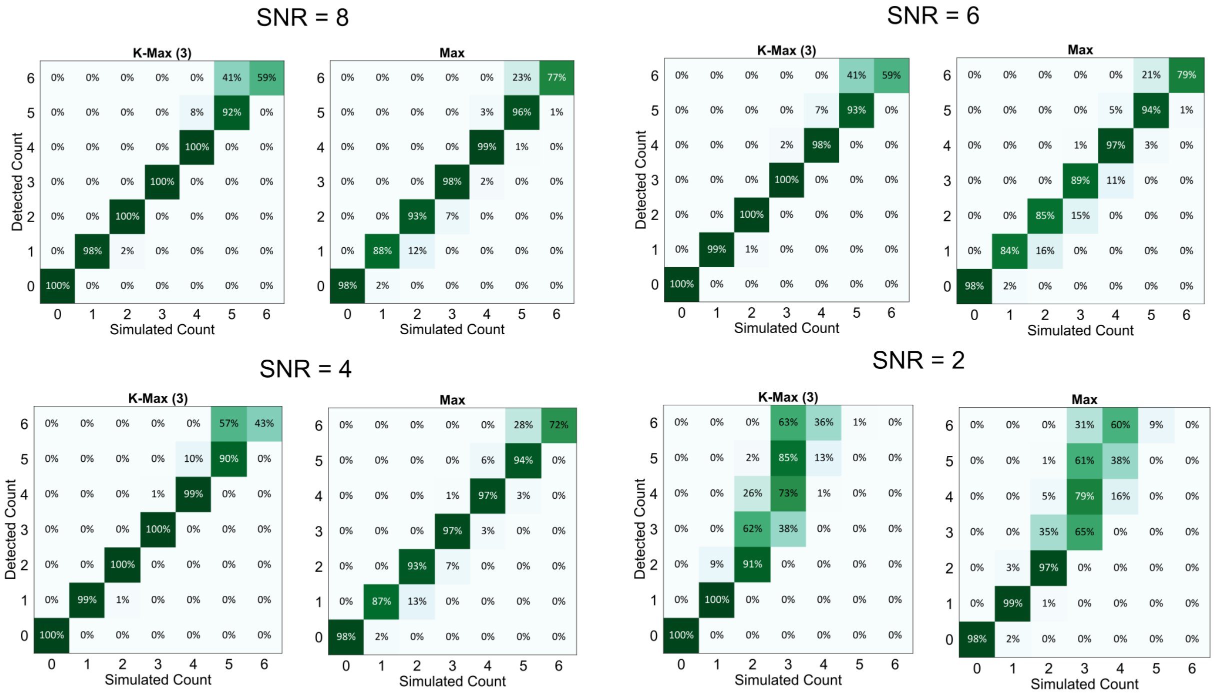


Figure S9: Confusion matrices showing the accuracy of the automated counting on simulated mRNA videos for SNRs of 8 (top left), 6 (top right), 4 (bottom left), and 2 (bottom right) using counting statistics of K-Max with $k$=3 and overall maximum.

#### S6.3 Effect of intensity adjustment

Due to the positional overlap of diffusing mRNA molecules in the confinement volume, the counting accuracy of mRNA in 5 μm microwells decreases significantly for copy numbers >4 (Fig. S3). To improve the counting accuracy for wells with >4 mRNA, we applied an intensity adjustment to the K-Max count for each experimental dataset. This adjustment was based on the concept that the summed intensity of all pixels in the masked well area (after denoising) should roughly scale with the number of labeled molecules in the well. This approach is similar to the method by which prior studies have estimated copy number (e.g., by scaling the encapsulated fluorescence intensity by that of a single labeled mRNA), but instead of only using measurements of single mRNA intensity, we leverage the high accuracy of the direct counting up to 4 mRNA (Section S2.3) for improved rescaling.

The intensity-based adjustment is applied as follows. In a given experimental dataset, we compute the sum of the pixel intensities within the masked well area over the first 10 frames and take the mean of this value. The use of only the first 10 frames minimizes the impacts of potential photobleaching. We apply this calculation for all wells with a detected number of mRNA molecules between 1 and 4. For these wells, we then model the summed well intensity as a function of the detected count and fit a linear scaling model. This model allows us to generate a prediction for the expected number of mRNA in a well based on the measured intensity over the whole well. We then apply this linear fit to the summed well intensities for all wells with ≥5 counts to better estimate the true mRNA copy number. A sample plot of the summed well intensities as a function of detected counts from 0 to 4 is given in Fig. S10. A Pearson correlation coefficient between 0.67-0.92 was obtained for all experimental datasets to which this recalibration fit was applied. Intensity-adjusted count values that were greater than the physical capacity of an LNP were truncated to 24 (n=2).

Given the moderate accuracy of the automated counting for wells with 5 detected counts (Fig. S9), the intensity adjustment was only applied to wells where K-Max = 5 if the summed intensity of the well exceeded a threshold given by:

$$I_{\text{well (5),i}} \geq 1.3 \times \overline{I_{\text{well (4)}}} \tag{S5}$$

where $I_{\text{well (5),i}}$ is the summed intensity of a given well with K-Max = 5 and $\overline{I_{\text{well (4)}}}$ is the

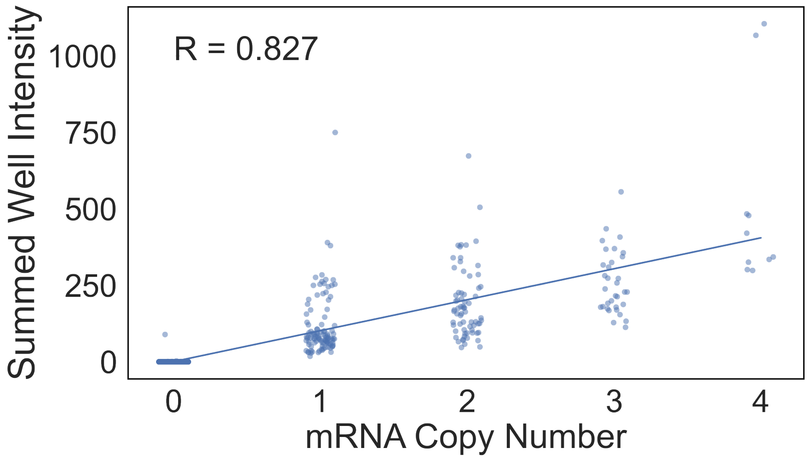


Figure S10: Sample scatter plot of the sum of pixel intensities within the well area as a function of the detected copy number for said well. The solid line shows a linear fit of the data.

mean summed intensity of all wells with K-Max = 4 in the same experimental dataset. This threshold was determined empirically through manual inspection. A specific linear fit was created for each experimental dataset to account for variations in RiboGreen fluorescence intensity due to incomplete staining across the array and small fluctuations in laser power and fluorescence background.

We compared this approach to the use of single mRNA intensities to recalibrate copy number as in prior studies.[3,24,29] Briefly, wells containing a single mRNA per K-Max were pooled and the intensity of each was computed as $I_0 \pi \sigma^2$, where $I_0$ is the amplitude and $\sigma$ is the width of a 2D Gaussian fit. For all wells with a detected count $\geq$5, the count was recalibrated by dividing the summed well intensity by the mean single mRNA intensity. Using simulations, we show that intensity adjustment using the sum of well intensities leads to stronger agreement between the true count and detected count than the approach of K-Max or using a single RNA intensity recalibration (Fig. S12).

We also applied the intensity-adjusted counting approach to simulated wells with counts less than 5 to further assess the validity of this approach. After applying the intensity adjustment, the distribution of detected counts for each corresponding simulated count became polydisperse, particularly for higher copy numbers (Fig. S11). Despite this polydispersity, the mean detected count for each simulated copy number, as shown in the rightmost column of Fig. S11, was highly accurate to the true value (i.e., within 5%). This indicates that while the intensity rescaling likely introduces uncertainty in the count estimate when applied to high copy numbers, the adjustment we apply provides an accurate reflection of the true count distribution on average.

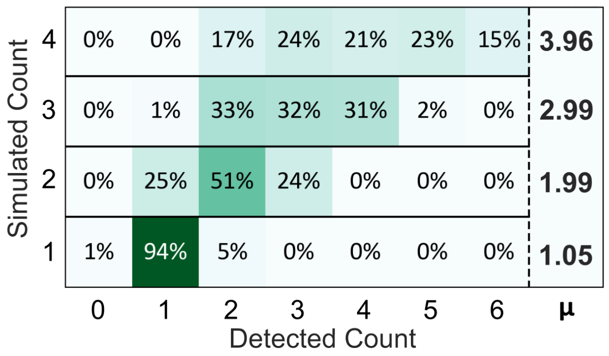


Figure S11: Confusion matrix showing the accuracy of intensity-based counting for well videos simulated with 1 to 4 particles (n=300 per row). The mean detected count for each simulated copy number is shown in the rightmost column and lies within 5% of the true count for all copy numbers.

#### S6.4 Effect of noise background

Simulated videos without particle signals were generated with varying levels of Poisson noise to test the software's sensitivity/false detection rate. For videos with a standard deviation of $1 \pm 0.1$ among pixel intensities, the average noise background observed in the experimental data, the false detection rate among 150 samples was 0. For videos with a standard deviation of $2 \pm 0.1$ or $3 \pm 0.1$ among pixel intensities, the false detection rate increased to 1%. The false positive rate was also tested as a function of the size of FOV. For a standard deviation of $1 \pm 0.1$ among pixel intensities, the following widths of FOV were tested: 40 pixels (corresponds to cropped area for a 5 um well), 80 pixels (corresponds to cropped area for a 10 um well), and 120 pixels (corresponds to cropped area for a 20 um well). For the same background noise level, the false detection rate ranged from 0% to 2% for FOVs of 40, 80,

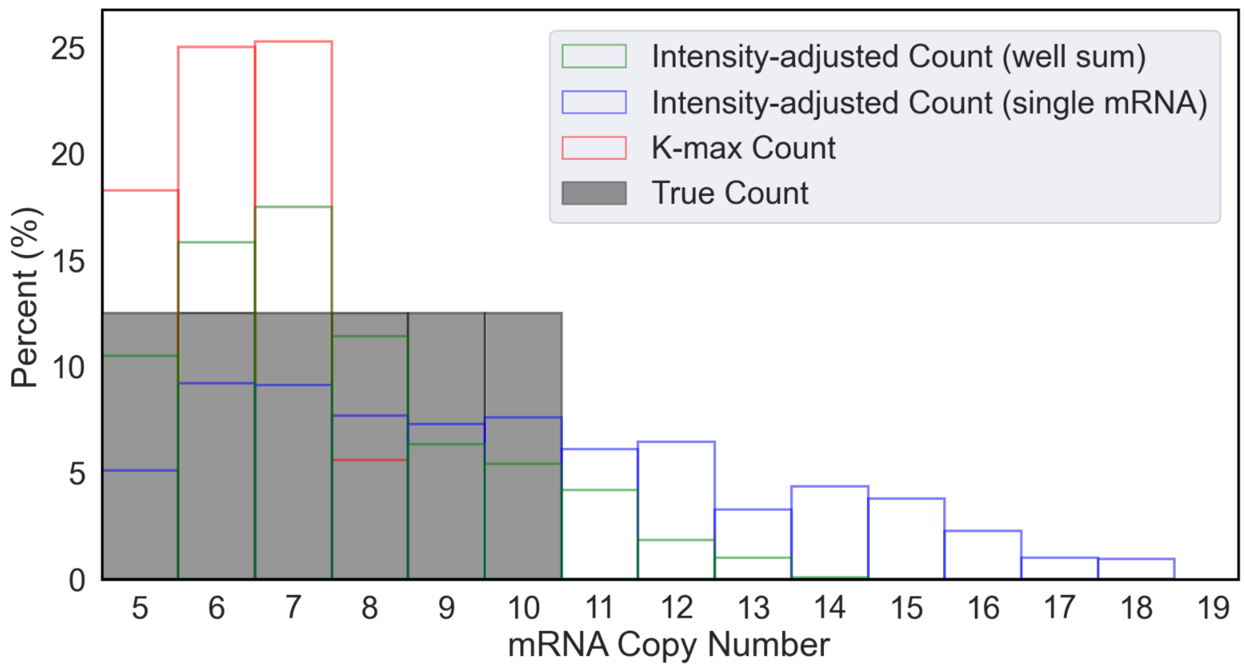


Figure S12: Distribution of mRNA counts as detected by K-Max (red), intensity adjustment by well sum (green), and intensity adjustment by single mRNA intensity (blue), all in comparison to the ground truth counts (grey).

and 120 pixels, indicating that for ideal counting accuracy in the case of larger wells, a new CNN should be trained with input images of comparable scale to decrease false detections.

##### S6.5 Effect of well size

This study is limited by the microwell geometries of the available CLiC flow-cells, where all available microwell arrays had a diameter of approximately 5 µm. The method of directly counting individual mRNA molecules relies on the separation of intensity peaks in the x-y plane of the confinement volume thus we wanted to investigate the specific effects of well size on the counting accuracy. A series of confusion matrices was generated to show the accuracy of the counting as a function of well size (5 µm vs 10 µm vs 20 µm) (Fig. S13) to demonstrate that this method could be used to count higher payloads provided different microwell geometries. For the standard K-Max statistic, overcounting is likely as the image area increases due to the additional noise pixels, as described in Section S6.4. To account for this, a dynamic statistic that scales with approximate the number of mRNA per well can be used. For this dynamic statistic, the overall median count would be used for approximate counts below 5 and the K-Max would be used for approximate counts above 5, where the K-Max is defined as the median of the 6 highest detections. Using this approach, up to 8 particles can be counted with ≥91% accuracy in 20 μm wells. This demonstrates that the mRNA counting method is adaptable to counting higher mRNA-LNPs payloads using larger microwells.

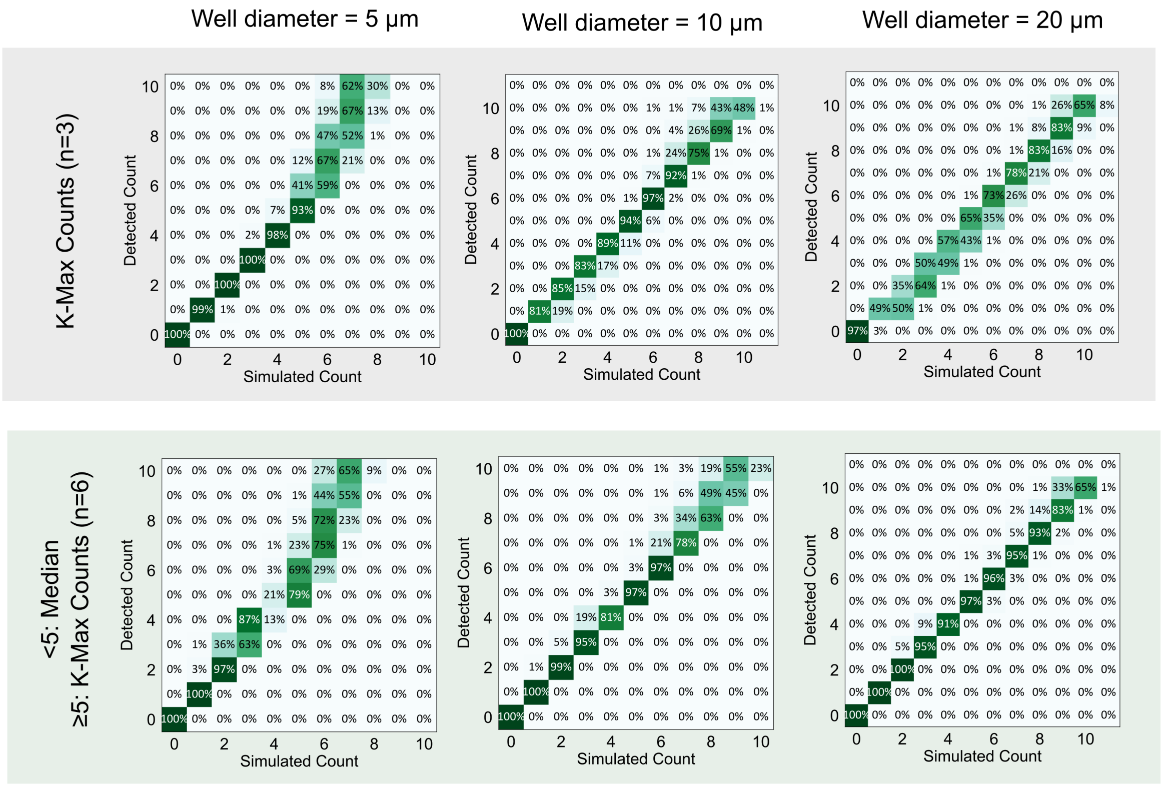


Figure S13: Confusion matrices showing the accuracy of the automated counting for microwell diameters of 5 μm (first column), 10 μm (second column), and 20 μm (third column). The standard K-Max with $k$=3 (top row) is compared to a modified K-Max that uses the overall median at low counts (<5) to prevent overcounting due additional noise pixels in the increased imaging area.

#### S6.6 Effect of exposure time

A series of confusion matrices was generated to show the accuracy of the automated mRNA counting as a function of exposure time (Fig. S14). This series was generated for various mRNA counts at an SNR of 6 to validate the design of the experimental imaging protocol. This series shows that an exposure time of 40 ms does not improve counting substantially compared to 20 ms and the additional motion blur also worsens counting at high copy number (Fig. S14, left). Conversely, an exposure time of 10 ms slightly worsens detection as compared to 20 ms for counts above 4 (Fig. S14, right). Therefore, a 20 ms exposure time is appropriate for accurate mRNA counting with the other imaging settings of the experiment taken into account.

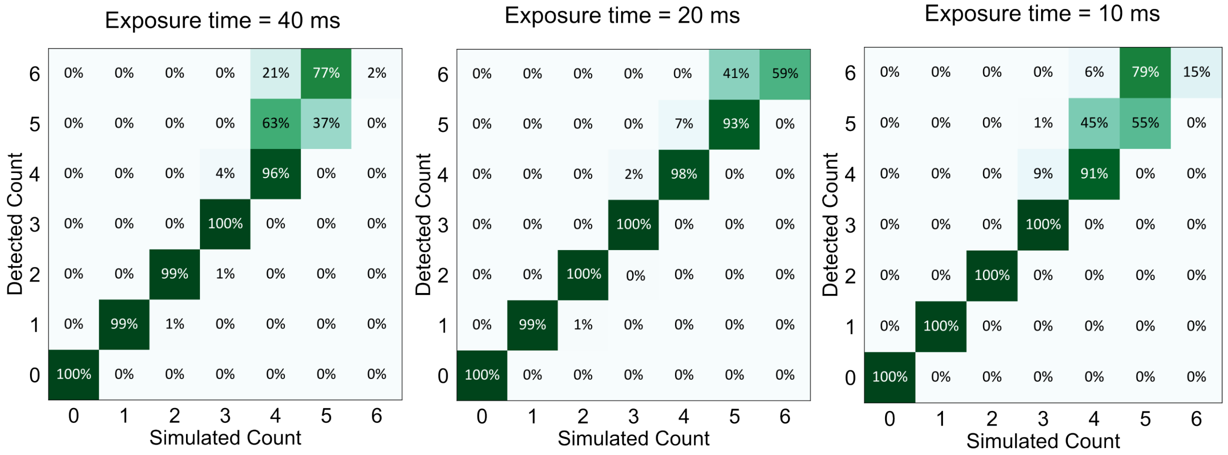


Figure S14: Confusion matrices showing the accuracy of the automated counting for well videos simulated with exposure times of 40 ms (left), 20 ms (middle), and 10 ms (right). All videos were simulated with a SNR of 6.

#### S6.7 Effect of number of frames

A series of confusion matrices was generated to investigate counting accuracy as a function of the number of frames (Fig. S15), assuming an exposure time of 20 ms, to validate the design of the experimental imaging protocol. Acquisitions with only 15 frames at an SNR of 4 result in worsened undercounting at high copy number due to an increased fraction of the frames having high positional overlap (Fig. S15, left). Acquisitions with 100 frames show negligible improvement in counting accuracy for double the imaging time (Fig. S15, right), thus 50 frames is a suitable acquisition length.

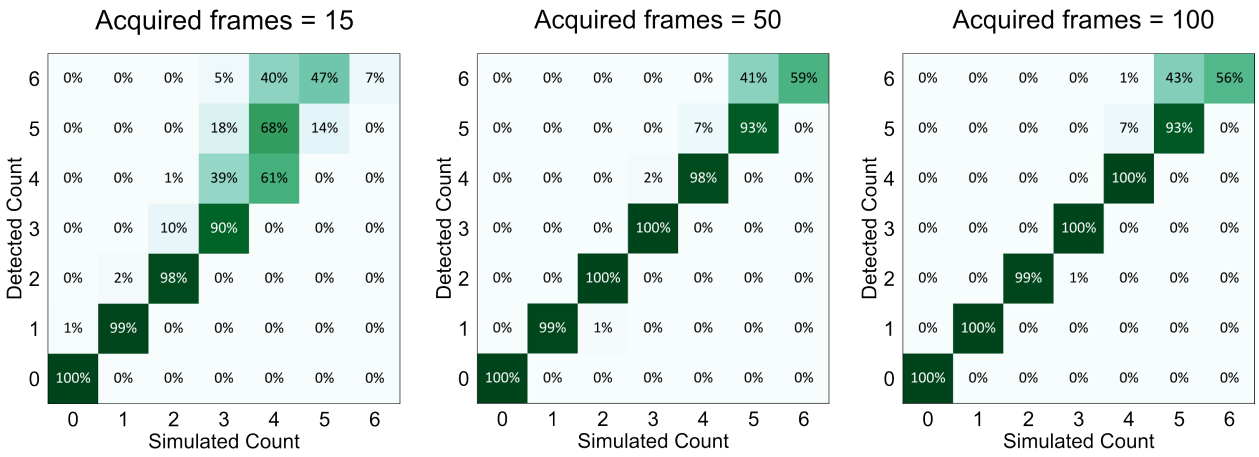


Figure S15: Confusion matrices showing the accuracy of the automated counting for well videos with 15 frames (left), 50 frames (middle), or 100 frames (right). All videos were simulated with a SNR of 6 and an exposure time of 20 ms.

(22) Kameyama, A.; Kaneda, Y.; Yamanaka, H.; Yoshimine, H.; Narimatsu, H.; Shinohara, Y. Detection of oligosaccharides labeled with cyanine dyes using matrix-assisted

laser desorption/ionization mass spectrometry. *Analytical chemistry* **2004**, *76*, 4537–4542.

(29) Li, S.; Hu, Y.; Li, A.; Lin, J.; Hsieh, K.; Schneiderman, Z.; Zhang, P.; Zhu, Y.; Qiu, C.;

Kokkoli, E.; Wang, T.-H.; Mao, H.-Q. Payload distribution and capacity of mRNA lipid nanoparticles. *Nature Communications* **2022**, *13*, 5561.
